# Inhibition stabilization is a widespread property of cortical networks

**DOI:** 10.1101/656710

**Authors:** A. Sanzeni, B. Akitake, H. C. Goldbach, C. E. Leedy, N. Brunel, M. H. Histed

## Abstract

Many cortical network models use recurrent coupling strong enough to require inhibition for stabilization. Yet it has been experimentally unclear whether inhibition-stabilized network (ISN) models describe cortical function well across areas and states. Here we test several ISN predictions, including the counterintuitive (paradoxical) suppression of inhibitory firing in response to optogenetic inhibitory stimulation. We find clear evidence for ISN operation in mouse visual, somatosensory, and motor cortex. Simple two-population ISN models describe the data well and let us quantify coupling strength. Though some models predict a non-ISN to ISN transition with increasingly strong sensory stimuli, we find ISN effects without sensory stimulation and even during light anesthesia. Additionally, average paradoxical effects result only with transgenic, not viral, opsin expression in parvalbumin (PV)-positive neurons; theory and expression data show this is consistent with ISN operation. Taken together, these results show strong coupling and inhibition stabilization are common features of cortex.

## I. INTRODUCTION

Extensive recurrent connectivity between nearby neurons is an ubiquitous feature of the cerebral cortex [1; 2; 3; 4]. Theoretical work has shown that the strength of recurrent coupling has a major impact on several computational properties of networks of excitatory (E) and inhibitory (I) neurons, including the speed of the response to external stimuli [5; 6], the ability of a network to sustain persistent activity [7], the capacity and robustness of memory storage [8], and the amplification of various input modes [9].

Strong excitatory recurrent coupling, however, can lead to unstable dynamics unless stabilized by inhibition. When recurrent connectivity is weak, excitatory cells can show stable firing rates independent of the activity of inhibitory cells. In networks with strong recurrent connections, excitatory-to-excitatory (E-E) connections amplify responses so that the excitatory network is unstable if the firing rates of inhibitory neurons are kept fixed. Stable excitatory network operation across a range of firing rates can be restored if inhibitory recurrent connections are sufficiently strong, allowing inhibition to track and balance excitation [5; 6; 7; 10; 11]. Such network models, with strong recurrent connections rendering the excitatory cells self-amplifying and thus unstable, and requiring inhibition for stability, are called inhibitory-stabilized networks (ISNs) [12].

Whether cortical networks function in the ISN regime, in which conditions they do so, and which cortical areas may operate as ISNs has been the subject of debate. Cat primary visual cortex (V1) shows behavior consistent with the ISN regime [12]. But since that work used sensory stimuli, it could not determine whether cat V1 operates as an ISN in the absence of visual stimuli (i.e. at rest). Based on these data and others, Miller and co-authors later developed a model called the stabilized supralinear network (SSN) [13; 14], which explains cat and ferret V1 responses to visual sensory stimuli of different sizes and intensities. The SSN shows ISN-regime operation for strong visual stimuli, but predicts that as sensory stimuli decrease in strength, the network should transition into a non-ISN state. Thus, an important open question has been whether cortical regions operate as an ISN even without sensory stimulation.

Optogenetic stimulation can give insight into ISN operation even in the absence of sensory stimuli. A prominent prediction of ISN models is *paradoxical inhibitory suppression*[10] — a counterintuitive decrease of inhibitory firing rate as inhibitory cells receive direct input (Fig. 1b). This paradoxical effect is a result of the strong synaptic coupling between excitatory and inhibitory cells. In an ISN, due to the strong recurrent coupling, more recurrent excitation is withdrawn after inhibitory stimulation than the stimulation adds, leading to a net decrease in input to stimulated inhibitory cells (see Fig. S1). This decrease in recurrent excitation is the cause of the paradoxical effect.

**FIG. 1:**
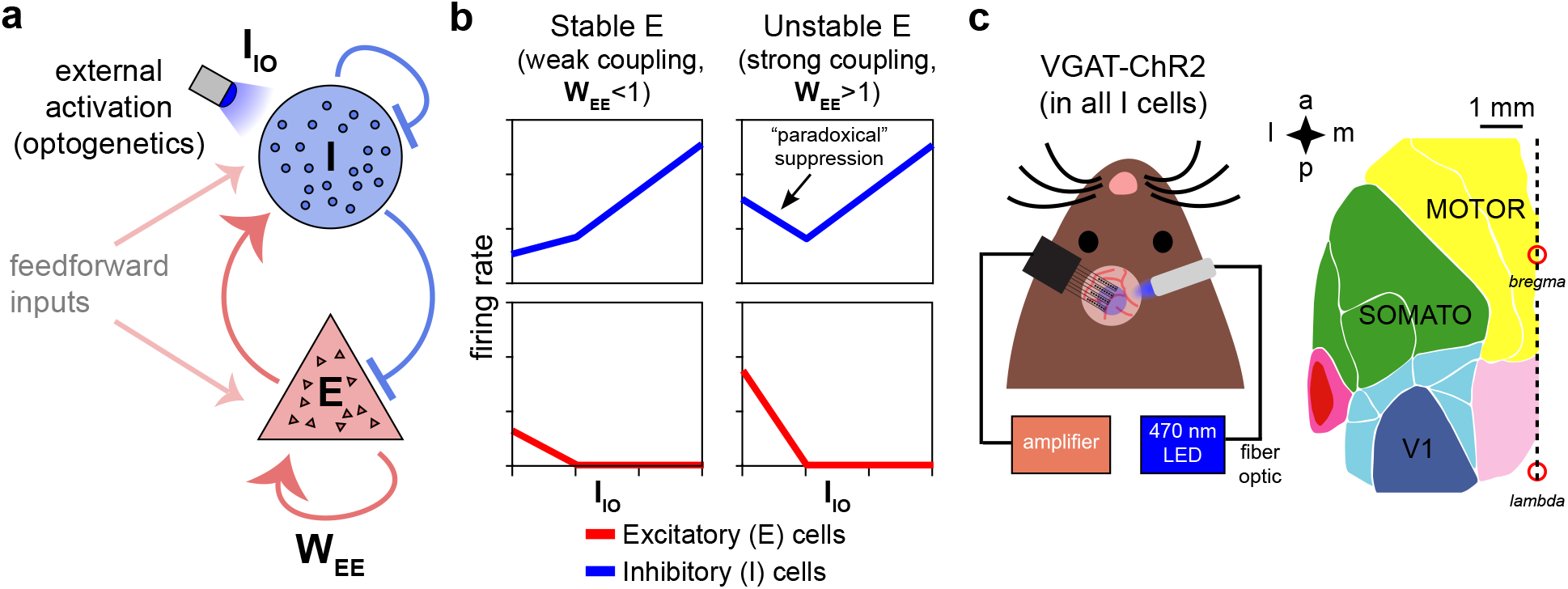
Model predictions of excitatory and inhibitory responses to inhibitory stimulation. (**a**) Schematic of model, showing connections between excitatory (E) and inhibitory (I) neuron populations. (*W_EE_*, a measure of the strength of E-E connections, is the key parameter controlling non-ISN vs ISN operation, see model discussion in Results.) (**b**) Predictions for average neural responses with weak recurrent coupling (left) and strong coupling (right), when inhibitory cells are externally stimulated (I_IO_). (**c**) Schematic of experiment. Extracellular recordings made in visual (V1), primary somatosensory, and motor/premotor cortices (see Fig. S2 for electrode locations; a: anterior, p: posterior, m: medial, l: lateral) while optogenetically stimulating inhibitory cells at the recording site in awake VGAT-ChR2-YFP animals.

Optogenetic ISN studies, largely performed in the mouse, have produced conflicting data about ISN operation. While some paradoxical changes in inhibitory currents have been observed in mouse primary auditory cortex (A1) [15; 16], several studies have found non-paradoxical effects in inhibitory firing rate or intracellular currents in mouse V1, A1, and primary somatosensory cortex (S1) [16; 17; 18], or found circuits with feedforward inhibition between two inhibitory cell classes that could produce paradoxical effects in a non-ISN [16]. Possible explanations for these varying observations include differences between anesthetized and awake states, differences between measurements of firing and intracellular currents, or, as we show experimentally below, differences in which, or how many, inhibitory cells are stimulated [18; 19; 20]. Additionally, in mouse V1, one study [21] found that excitatory and inhibitory currents scale as predicted in the SSN with increasing visual stimulation. But this study of SSN phenomena also leaves open the question whether mouse V1 operates as an ISN without sensory stimuli (see also [19], their Discussion).

Here, we examine whether cortical networks operate as ISNs (Fig. 1) at rest, by combining optogenetic stimulation of inhibitory cells, extracellular recordings in awake animals from several cortical areas, and theoretical analyses. Using a mouse line that allows stimulation of all inhibitory cells together, we find clear evidence for ISN operation even without sensory stimulation in multiple mouse cortical areas and both in superficial and deep layers. By providing multiple experimental and theoretical lines of evidence for ISN operation, our results argue against non-ISN explanations (e.g. against paradoxical suppression arising from one inhibitory population disinhibiting another in the absence of strong recurrent coupling).

We also address whether stimulating a single genetic subclass of inhibitory neurons in V1, the parvalbumin-positive (PV) cells, can produce paradoxical inhibitory effects. We find differences in paradoxical responses between viral and transgenic expression strategies. These differences are explained by an ISN model where the fraction of stimulated PV cells is varied [18; 20], and are supported by histological measurements, supporting the idea that cortical responses to inhibitory input can be highly dependent on the number of stimulated cells [20].

Together, these results support the idea that the cortex operates in the ISN regime across a range of areas and brain states.

## II. RESULTS

### Mouse primary visual cortex is inhibition stabilized

We first describe experiments where all inhibitory neurons were optogenetically stimulated. We expressed an excitatory opsin (Channelrhodopsin-2; ChR2) in all inhibitory neurons using a transgenic mouse line (VGAT-ChR2 [22]). We delivered blue light to the surface of the cortex, while recording activity extra-cellularly with multi-site silicon recording arrays (Fig. 1c). Mice were given drops of reward approximately once per minute (Methods) to keep them awake and alert. Because neurons in the superficial cortical layers showed the largest responses to the stimulation light (Fig. S3), we restricted our analyses to units within ≈400 *μ*m of the cortical surface (Methods), which primarily includes neurons in layer 2/3 and upper layer 4 [23]. We sorted units into single- (separated from noise and other units) and multi-units (lower SNR or containing several apparent units) by waveform (Methods; using a more restrictive threshold for unit inclusion does not affect our conclusions, Fig. S5), and we excluded the multi-units from further analyses.

The majority of recorded units were suppressed by light (146/167 single units, 87%; Fig. 2b-d; 4 animals, 7 recording days). Given that approximately 80% of cortical neurons are excitatory [24; 25], and that extracellular recordings can show several biases in sampling cortical neurons, [26; 27], this measured percentage could be consistent with either the presence or absence of paradoxical suppression in inhibitory neurons. Therefore, we classified recorded units as inhibitory or excitatory using *in vivo* pharmacology. (Our results are confirmed by other classification methods: identifying inhibitory neurons using their response at high laser intensity or by waveform width give similar results, see below and Fig. S8.)

**FIG. 2:**
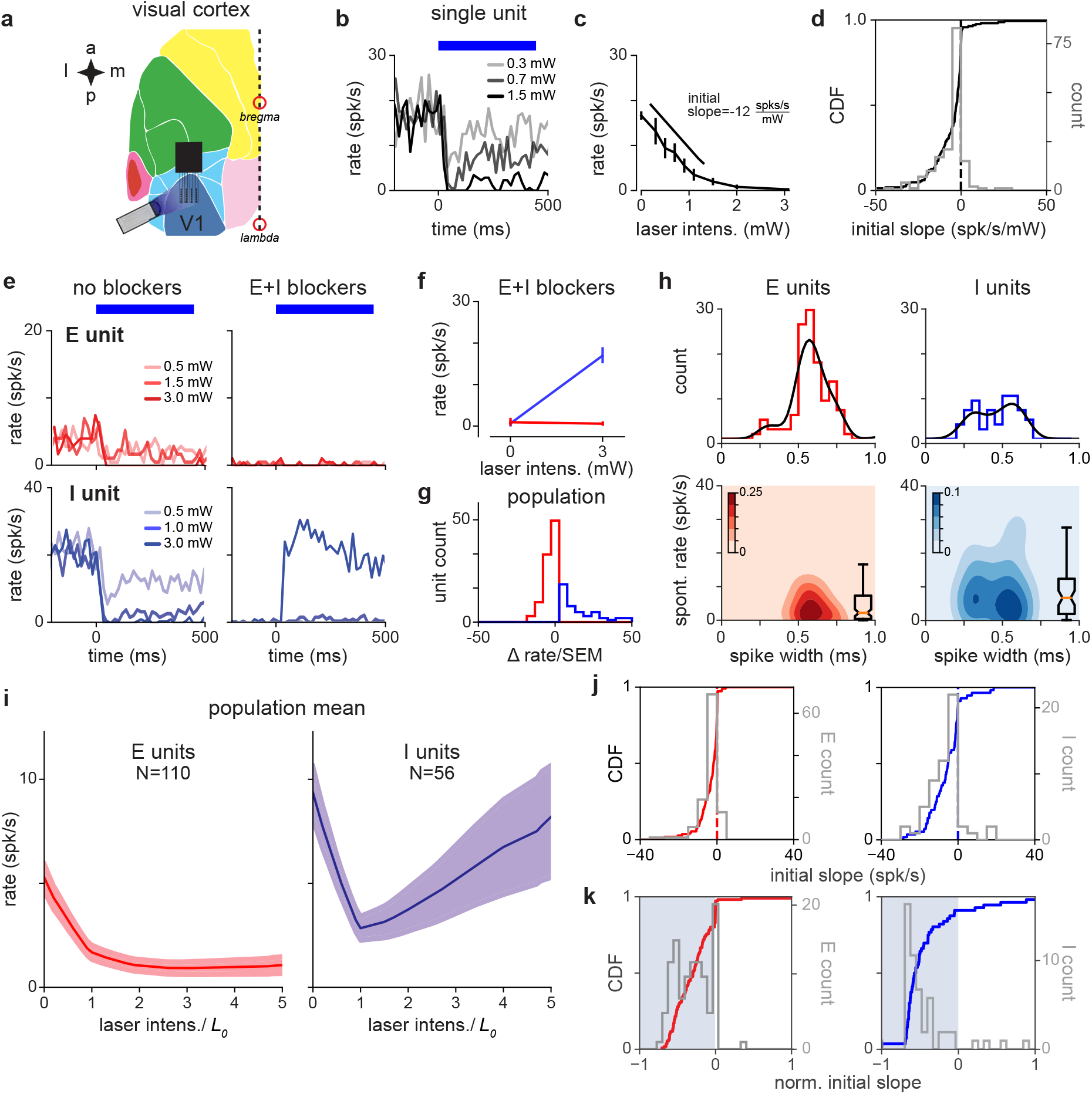
V1 response is consistent with inhibition stabilization. (**a**) Schematic of experiment, showing recordings/stimulation in V1. a: anterior, p: posterior, l: lateral, m: medial. V1 recording locations and intrinsic signal imaging to define V1 shown in Fig. (S2). (**b**) Example recorded unit; greater laser power produces greater firing rate suppression. Blue bar: duration of constant-power laser pulse. (**c**) Average firing rate of example recorded unit (rate is steady-state firing rate, over last 300 ms of laser stimulus). Suppression is quantified via initial slope of firing rate vs laser intensity (Methods). (**d**) Distribution of initial slopes for all recorded units. (**e**) Example cell responses without (left) and with (right) blockers. Top: Cell classified as excitatory; does not respond to laser in presence of blockers (CNQX, APV, bicuculline; Methods). Bottom: Cell classified as inhibitory; increases firing rate to laser in presence of blockers. (**f**) Firing rates for units in (e) in presence of blockers; inhibitory cell (blue) shows a large positive change. (**g**) Distribution of firing rate changes to laser stimulus with blockers. Red: E-classified units; blue: I-classified units. (**h**) Spontaneous firing rates and waveform width (time from waveform first local minimum to first local maximum) for units classified as E and I via the method in panel g. (**i**) Average population response for units classified as E and I. inhibitory cells show a prominent paradoxical effect. (To account for variations in tissue optical properties across recording days, we combined data by fitting a piecewise-linear function to the average inhibitory response for each day to find a change in slope (*L*_0_: always found to be a minimum between 0.3 and 2.7 mW, see Fig. S4) and rescaling the laser intensity so *L*_0_ = 1.) (**j**) Distribution of initial slopes for units classified as E (red, left) and I (blue, right). (**k**) Distribution of normalized initial slopes 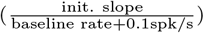, to more clearly show small slopes, for units classified as E and I. Several excitatory cells with very low firing rates in the no-blocker case have near-zero normalized slopes. Errorbars: ±1 SEM (as throughout figures, unless otherwise noted.)

We first recorded neurons’ responses to optogenetic stimulation, and then applied blockers of excitatory and inhibitory synapses (CNQX, AP5, bicuculline; which block AMPA, kainate, NMDA, and GABA-A synapses; Methods). When recurrent synapses are substantially suppressed, inhibitory cells expressing ChR2 should increase their firing rate to optogenetic input, while excitatory cells’ firing should not increase. Indeed, by measuring responses to inhibitory stimulation in the presence of E and I blockers, we were able to classify units into two groups (Fig. 2e). Units in the first class (putative excitatory) were silent for strong stimulation and units in the second class (putative inhibitory) increased their firing rate in response to stimulation. To quantitatively classify cells, we measured the change in activity induced by light when the blockers had been applied (Fig. 2f-g); cells were labeled inhibitory (33%; 56/167 units) if the change compared with baseline produced at maximum laser intensity was positive (according to Welch’s t-test, *p* < 0.01), and excitatory otherwise. Choosing different classification thresholds did not affect our results (Supp Fig. S7), examining a subset of cells with the most stable waveforms over time did not affect our results (Fig. S6), and, finally, examining a subset of inhibitory units where waveforms had high signal-to-noise ratio also did not qualitatively change the results (Fig. S5). Therefore, we used these response differences in the presence of blockers to identify recorded inhibitory units.

Pharmacologically-identified inhibitory and excitatory units showed differences in waveform width and spontaneous firing rate (Fig. 2h), though these two factors were insufficient alone to completely classify inhibitory cells (i.e. insufficient to predict whether units would increase firing rate in the presence of blockers). Identified excitatory cells had a wider waveform (0.58±0.01ms) and lower firing rate (6.0±0.8 spk/s) than inhibitory cells (width 0.46±0.02 ms, rate 10.7±1.7 spk/s). However, inhibitory neurons’ width was broadly distributed, supporting the idea that that some inhibitory classes can have broader waveforms than others [28; 29].

Once recorded units were classified as excitatory or inhibitory, we could ask whether the same units before adding blockers (i.e. with the animal in the awake state, with recurrent connections intact) showed paradoxical inhibitory effects. Indeed, inhibitory neurons showed strong suppression when stimulated (Fig. 2i-j). This paradoxical inhibitory suppression is one piece of evidence that an ISN model is a good description of the upper layers of primary visual cortex. (Below we show that response dynamics and responses at high laser intensity provide additional evidence for an ISN.)

These experiments were done without sensory stimulation, as animals viewed an unchanging neutral gray screen. Because visual cortical neurons’ firing rates increase with increasing contrast across a range of overall luminance levels, we expected that we would see little difference between data collected with a neutral gray screen and data collected in the dark. Confirming that expectation, in one experimental session we measured inhibitory responses to stimulation while animals were in the dark (with the blue optogenetic light shielded with light-blocking materials), and found paradoxical suppression (N=6/6 inhibitory units with initial negative slope, median initial slope −6.9 ±3.5 spk/s; p=0.046, median < 0, Mann-Whitney test).

In an ISN, paradoxical inhibitory suppression arises because exciting inhibitory cells in turn suppresses excitatory cells, which withdraw excitation from the stimulated inhibitory cells. Then, because more excitatory input from recurrent sources is withdrawn from inhibitory cells than excitatory input from optogenetic stimulation is added, the net steady-state response of inhibitory cells is suppression. For this mechanism to hold, inhibitory cells must transiently increase their firing, even if only a small number of inhibitory spikes are generated, before the paradoxical suppression [10; 12]. Therefore, we looked for transient increases in inhibitory firing with short latencies after stimulation, and found a small increase in inhibitory firing rate immediately after stimulation, before the larger paradoxical suppression (Fig. 3). The transient was brief (FWHM 7.2 ms), and so the actual number of extra spikes fired in the transient was small: only 0.03 extra spikes were fired on average per inhibitory-classified unit per stimulation pulse, though because we used 100 pulse repetitions, the majority of classified inhibitory units showed a detectable initial transient (Fig. S10).

**FIG. 3:**
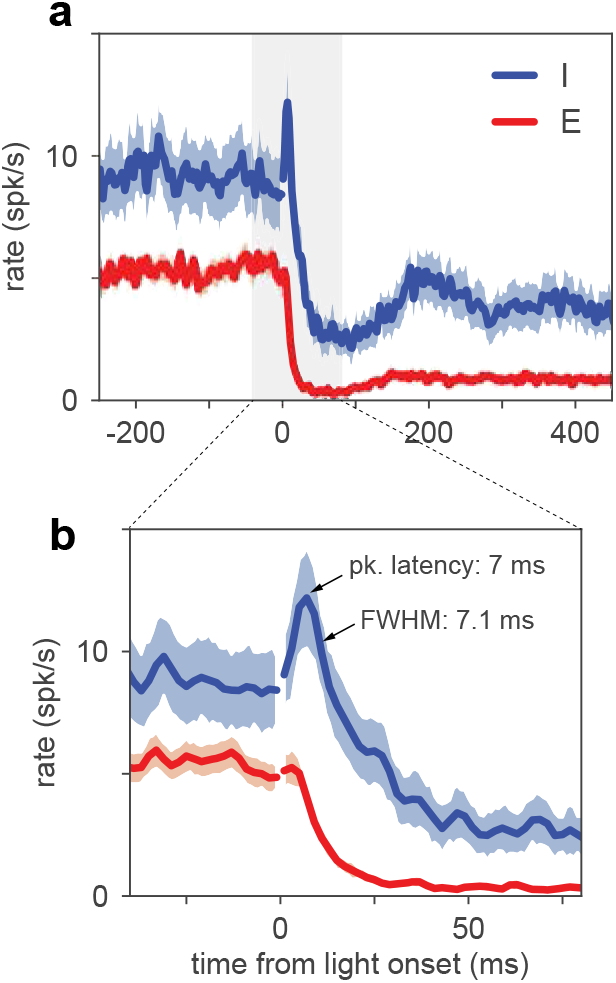
Inhibitory neurons show a small increase in firing before paradoxical suppression. (**a**) Average timecourse of neural responses from units classified as excitatory (N=110) and inhibitory (N=56). Light pulse is constant-intensity and lasts for 800 ms (high-intensity pulse, 2.6·*L*_0_; stimulation strength larger than *L*_0_ is predicted by ISN models to produce larger transients than at *L*_0_, though steady-state I rates are larger than the minimum; see also Fig. S10). Data from VGAT-ChR2 animals. Heavy lines: population mean rate, smoothed with a LOWESS filter. Shaded regions: SEM (Methods). Note that the onset of the light pulse produces an artifact in the 0-ms time bin only; we drop spikes at that time point (broken lines, time=0) to remove the artifact. Inhibitory neurons show a brief increase in firing before suppression, and the increase above baseline is statistically significant (inhibitory firing rates over 12 ms window after pulse are greater than a matched-duration interval before pulse, *p* < 0.01, KS test). (**b**) Same data as (a), enlarged to show the initial transient (time range here is indicated in (a) by gray shaded region). Inhibitory initial positive transient has peak amplitude 3.6 spk/s above baseline, latency to peak 7 ms, full width at half maximum 7.1 ms, and inhibitory rate crosses baseline into suppression at 13.1 ms.

Another ISN prediction is that inhibitory cells should increase their firing rates at high laser intensities. In a recurrent network operating as an ISN, stimulating inhibitory cells first decreases inhibitory firing rates, but stronger stimulation eventually suppresses excitatory cells until they reach a firing rate where the excitatory network is stable without reactive inhibition. Then, increasing stimulation of inhibitory cells beyond this point produces increases in inhibitory firing rates, as the network moves into a non-ISN regime (Fig. 1C). Indeed, we observed this in our excitatory and inhibitory identified populations (Fig. 2H).

While our extracellular recording approach has advantages, such as allowing precise measurements of spike rate dynamics, and recording a large number of E and I neurons in the same experiment, one concern might be that misclassification of excitatory waveforms into inhibitory units could mask inhibitory cell responses. However, confirming that our recording procedures accurately measured inhibitory waveforms, (1) paradoxical suppression is maintained when we make our classification thresholds more stringent (restricting analysis to units with highest waveform signal-to-noise ratio, Fig. S5; varying classification threshold in the presence of blockers; Supp. Fig. S7), (2) units classified by spike width, or by response at high laser power, also showed clear paradoxical effects (Fig. S8), and (3) inhibitory, but not excitatory, response dynamics showed an initial increase followed by suppression (Fig. 3), the response patterns expected for E and I neurons in an ISN (Fig. S10). Also, as shown below, in parvalbumin-positive neuron stimulation with viral transfection, using the same waveform-sorting procedures, we observed no average paradoxical effects. Together, these observervations argue that paradoxical effects in VGAT-ChR2 animals do not arise due to waveform sorting, but instead due to the properties of the cortical inhibitory network.

Taken together, these results show that mouse V1 responses are consistent with those of a strongly coupled network whose activity is stabilized by inhibition.

### Model-based inference of network coupling strength and stability

We next used these data to infer network connection parameters in a standard two-population model that describes the dynamics of population-averaged firing rates of excitatory and inhibitory neurons (see Eqn. 6; Methods). This parameter inference is possible because we obtained E and I response measurements in three pharmacological conditions (no blockers; with E blockers; with E and I blockers; Fig. 4a), and with data from these three conditions, there are more experimental observations than model parameters (Methods: Model degrees of freedom). The model uses a rectified-linear single-neuron transfer function; using nonlinear transfer functions give similar results (Fig. S12 and Appendix A).

**FIG. 4:**
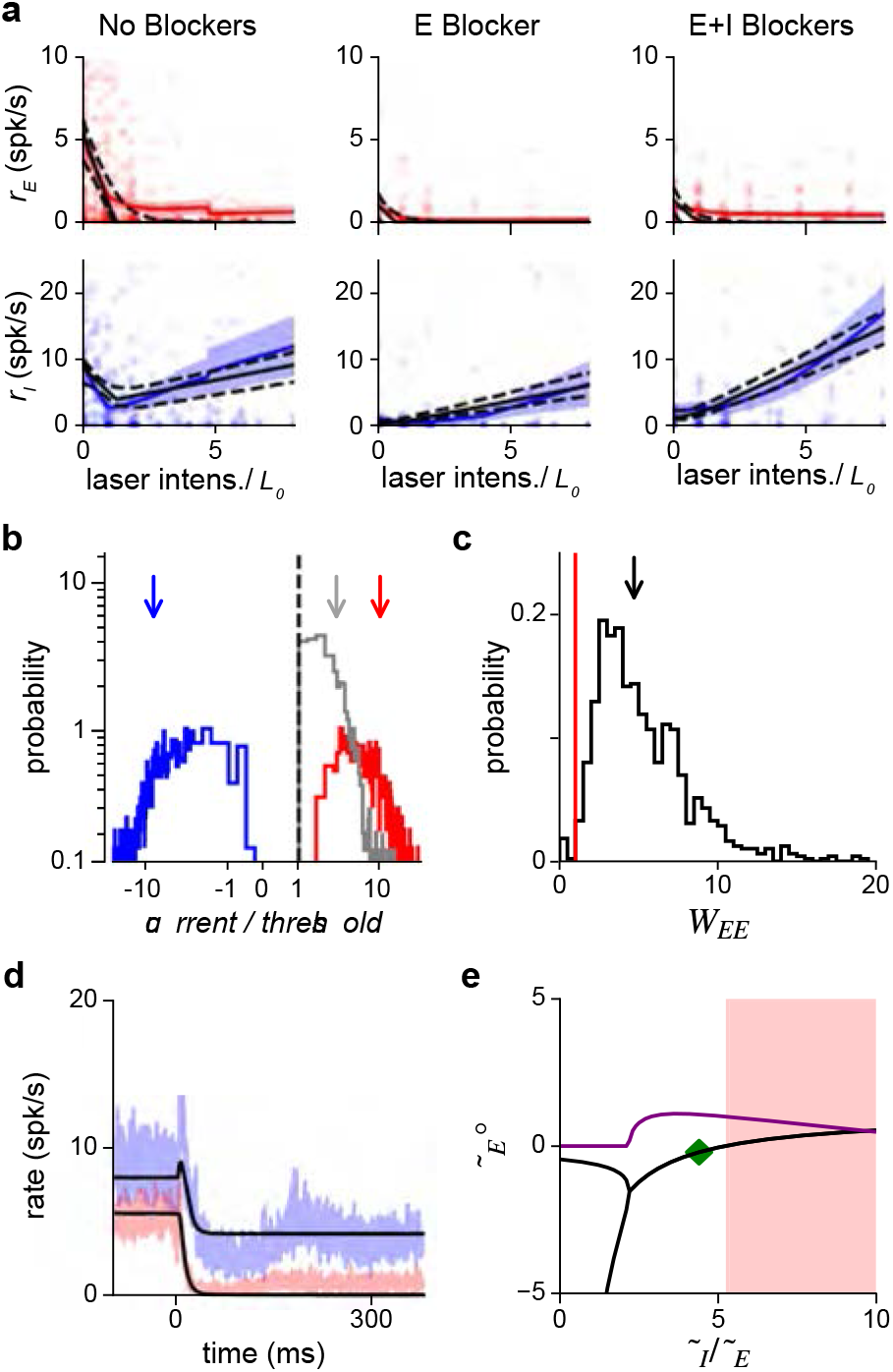
Population response is consistent with a network with moderately strong coupling. (**a**) Average excitatory (red) and inhibitory (blue) response measured in three conditions: without synaptic blockers, with only the excitatory blockers present, and with both E and I blockers present. Model (Eq. 6) that best fits data: black continuous line; dashed: 1 s.d. obtained by bootstrap; solid red/blue lines: E/I data means; shaded region: ±1 SEM. Because we applied blockers sequentially (generating separate E and E+I blocker measurements) the number of independent observations were increased, allowing the inference of all model parameters (Math. Methods; S11). Small steps in left panels (no blockers) arise because a subset of experimental days used a maximum laser intensity of 5 · *L*_0_. (**b**) Excitatory (red), inhibitory (blue), and net (gray) current influx into excitatory cells predicted by the model. Arrows: medians (over bootstrap repetitions); E: 10.9, I: −8.63; modes: E: 2.5, I: −2.7. (**c**) Distribution of *W_EE_* values compatible with the data; the red line represents the transition point between the ISN (*W_EE_* > 1) and the non-ISN (*W_EE_* < 1) regime. Median (arrow) 4.7; mode 2.5. (**d**) Estimation of time constants of E and I populations. Black line shows the dynamics resulting from fitting the data (blue: I population; red: E population; shaded region ±1 SEM) with the model for the same laser power shown in Fig. 3. Best-fit values are *τ_I_* = 34.3 ms and *τ_E_* =7.8 ms; note that network response dynamics shows faster time constants, due to recurrent network effects. The full model provides a good approximation to the dynamics even though it is constrained to simultaneously fit the time constants and the responses at different intensities. (**e**) Stability analysis. Real (black) and imaginary (purple) parts of the eigenvalues of the Jacobian matrix as a function of the ratio *τ_I_*/*τ_E_*. Imaginary part (purple) greater than zero signifies the network can show damped oscillations when being driven to a new stationary point. Note that these damped oscillations are not seen in (d) because of rectification. When the real part (black) is greater than zero, the network is unstable (shaded red area).

First, we find that the model, despite being overconstrained by the data, is a good description of V1 responses (Fig. 4a). The model also makes a number of predictions that are verified in the data. It predicts that the inhibitory firing rate slope for high laser intensities should be the same in the no-blocker and E-blocker conditions, as seen in the data (Fig. 4a, blue: left vs. middle columns). The model makes three other predictions: that the reversal point of inhibitory cells (*L*_0_, the point at which initial negative inhibitory slopes become positive), in all three experimental conditions, should match the point where excitatory cells change their slope to become nearly silent. These effects are also seen in the data (e.g. clearly visible in the no-blocker case, Fig. 4a, left column). Note that, although these predictions have been derived in a threshold-linear model (Methods), they are general features of network models which show a transition from non-ISN to ISN as the activity level of the excitatory population increases.

One prediction of balanced-state models with strong recurrent coupling [5] is that the total excitatory and inhibitory currents should each be large, though they cancel to produce a small net input. To check this, we computed from the model the currents flowing into excitatory cells (Fig. 4b). The excitatory and inhibitory currents are each approximately ten times larger than threshold (medians are 10.9 and −8.63, respectively), and their sum is small compared to the magnitudes of E and I currents (median net current/threshold = 2.2). The model also allows us to infer the value of the key parameter for inhibition stabilization, *W_EE_* (the excitatory self-amplification). The excitatory network is unstable, requiring inhibition for stabilization, and yielding an ISN, when *W_EE_* > 1. As expected when paradoxical effects are present, the fitted value is greater than 1 (Fig. 4c: median 4.7, mode 2.5). This value is not much larger than unity, suggesting recurrent coupling is not as strong as what the balanced-state model would predict [5; 6; 7] but is rather more consistent with a moderately-coupled network [13; 14; 30]. In Appendix A, we extend this model analysis to examine networks of spiking neurons which feature a fixed level of input noise, and find qualitatively similar results. The model shows that measured activity is consistent with the mean input to cells being below threshold. This implies the recorded cells operate in the fluctuation-driven regime, matching expectations from cortical response variability [5; 6; 7; 31; 32]. Moreover, when input fluctuations are taken into account in the model, mean excitatory and inhibitory currents are found to be of order of the distance between rest and threshold, in agreement with recent estimates suggesting the cortex operates in the loosely balanced regime [33].

Finally, we computed the parameter ranges over which the network is stable, as a function of the time constants of E and I firing rate dynamics, *τ_E_* and *τ_I_*. We find, by computing the eigenvalues of the connection matrix (Methods) that the network is stable for *τ_I_*/*τ_E_* in the interval [0, 5.3], and the observed dynamics are consistent with *τ_I_*/*τ_E_* = 4.4 and *τ_E_* = 7.8ms (Fig. 4e); that is, with excitatory and inhibitory synaptic time constants of the same order. The values of *τ*_(*E,I*)_ inferred from the response dynamics are within the stable region, providing further support that the ISN model is a good description of the effects we measure. (Note that the time constants of the opsin may increase these estimates, so the values of *τ_E_* and *τ_I_*, while small, are only upper bounds on the true dynamic response of the network to an instantaneous conductance change.) The ratio between E and I time constants is close to the point at which the network becomes unstable through an oscillatory instability (Fig. 4e). The fact that the inferred network model describes the data and is stable with *W_EE_* > 1 supports the experimental observation that the underlying cortical network is inhibition-stabilized.

### Inhibition stabilization is present in other cortical areas

The data above show that the superficial layers of mouse V1 are inhibition stabilized. Another open question is how general the ISN regime is across cortical areas. We performed experiments to look for signatures of inhibitory stabilization in two other cortical areas (Fig. 5): somatosensory (body-related primary somatosensory, largely medial to barrel cortex) and motor/premotor cortex. (See Fig. S2 for locations of all recording sites.) As in V1, we recorded units extracellularly, restricted analysis to the superficial layers of the cortex, and examined responses to optogenetic stimulation of all inhibitory cells (as above, using the VGAT-ChR2 line).

**FIG. 5:**
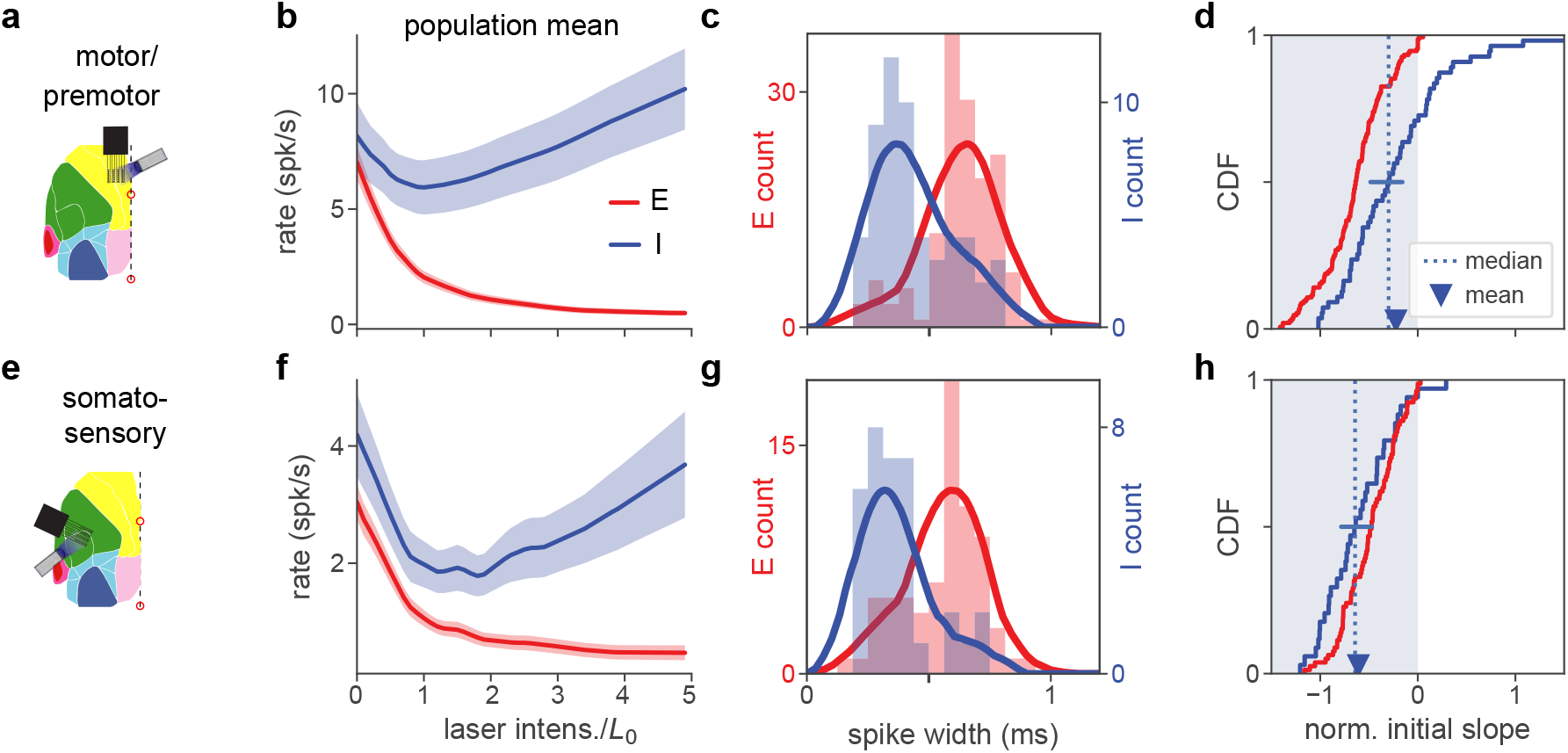
Inhibition stabilization across cortical areas. (**a**) Motor/premotor cortex recordings (see Fig. S2 for recording locations). (**b**) Motor cortex population firing rates for E and I units. Initial mean response of inhibitory cells is negative, showing paradoxical suppression. Mean rate is significantly reduced (*p* < 10^−4^, paired t-test, rate at 0 vs rate at *L*_0_). (**c**) Spike width distributions for E and I units. Units are classified as E or I here by response at high laser power (Methods), independently of spike width, which nonetheless varies with E or I unit identity. (**d**) Normalized initial slope distributions for all units. Red: E. Blue: I. Both mean and median of initial slopes are negative (paradoxical). Mean I slope is negative (*p* < 0.01, t-test). Horizontal bar at I median shows 95% confidence interval calculated by bootstrap. (**e-h**) Same as (a-d), but for recordings from somatosensory cortex. In (g), highest red bar is truncated for visual clarity (value is 22). Mean I rate (f) is sigificantly reduced (*p* < 10^−7^, paired t-test). Mean I slope (h) is negative (*p* < 10^−10^, t-test). Horizontal bar in (h) shows 95% CI around median slope.

In these experiments, we classified neurons as inhibitory based on their activity at high laser intensity. In our V1 experiments, paradoxical effects are clear whether neurons are classified in this way, or with the pharmacological method (Fig. S8). This high-laser-power classification is likely more stringent (i.e. may reject some inhibitory cells if they are suppressed by other inhibitory cells, due e.g. to heterogeneity of recurrent connections) than pharmacological classification. Supporting that this classification may be more stringent, some pharmacologically-classified inhibitory cells show paradoxical effects but little increase at high power (Fig. S9).

In both the recorded areas (Fig. 5), inhibitory neurons showed paradoxical effects, supporting that these cortical areas do also operate as an ISN. Similar to the V1 data, and as expected in an ISN, both areas showed a transition to a non-paradoxical response at large laser intensity. Compared to the somatosensory data, the motor/premotor recordings show a smaller average suppression (Fig. 5b) and more inhibitory units with non-paradoxical effects (rate increases with stimulation; Fig. 5d). However, in both areas, means and medians are significantly negative (see legend for statistical tests). Thus, the superficial layers of both sensory cortical areas we examined (visual and somatosensory), and one non-sensory area (motor/premotor) showed responses consistent with strong coupling and ISN operation.

### Differences in paradoxical suppression with viral or transgenic opsin expression can be explained by different numbers of stimulated cells

Up to this point, we have studied paradoxical suppression by stimulating an opsin expressed in all inhibitory cells via the VGAT-ChR2 mouse line. A remaining question is whether stimulation of a subclass of inhibitory neurons also yields paradoxical suppression. Even in an ISN, stimulation of any single subclass of inhibitory cells need not produce paradoxical suppression (see Methods and [14; 18; 19; 20]). To study this, we stimulated parvalbumin-positive (PV) neurons, which provide strong inhibitory input to other cells (Fig. 6). PV basket cells are the most numerous class of cortical inhibitory cells and make strong synapses near the somata of excitatory cells (for review, see [25]), and PV stimulation effectively suppresses network firing rates (e.g. [34; 35]).

**FIG. 6:**
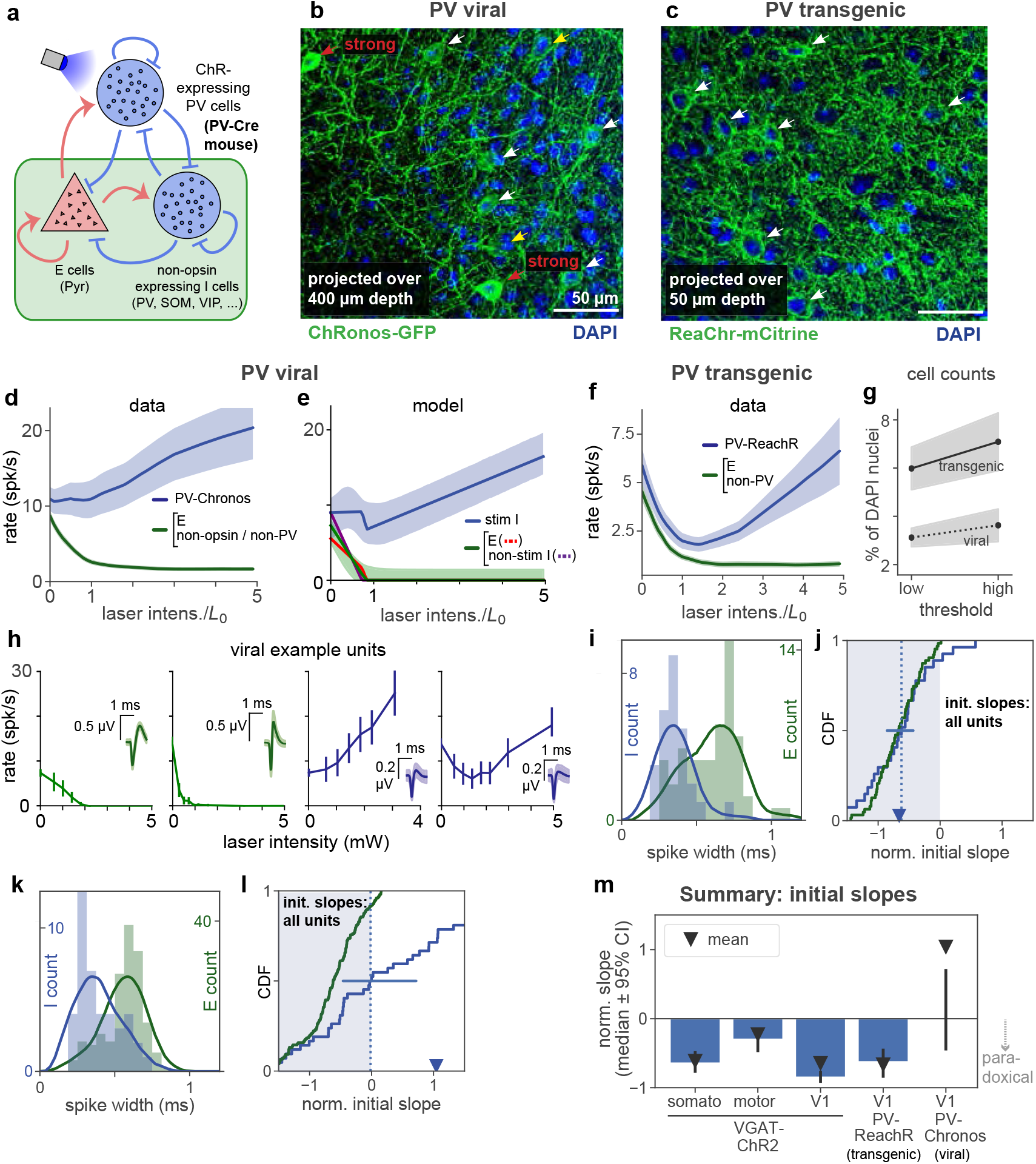
Stimulation of parvalbumin-positive inhibitory neurons shows paradoxical effects depend on number of cells stimulated. (**a**) Schematic of cell populations: opsin-expressing PV cells, non-stimulated inhibitory cells (non-PV inhibitory cells: SOM, VIP,…, and non-opsin PV cells), and pyramidal (E) cells. Data are from two experiments: Viral expression of opsin (Chronos) in PV neurons (AAV-Chronos;PV-Cre), and 2, transgenic expression of opsin (ReaChR) in most or all PV neurons (ReachR;PV-Cre). Responses shown are steady-state responses after light stimulation (Methods), to avoid differences in opsin kinetics affecting results. (**b,c**) Histological characterization of viral (b) and transgenic (c) expression in superficial layers. Viral expression shows more variability across neurons (cf red and white arrows), and fewer expressing cells (viral image is a projection across greater depth than transgenic). (**d**) Responses to stimulation with viral expression. Non-PV-Chronos (E, non-Chronos, or non-PV) cells: green (N=152). PV-Chronos (blue) cells (N=42, 21%), identified by responses at high laser power (Results; see Fig. S8 for validation against pharmacology-based classification in V1 data). Compared to when all inhibitory cells are stimulated (VGAT-ChR2 mouse line, Fig. 2), weaker paradoxical suppression is seen (blue line; mean rate is not significantly suppressed: *p* > 0.05, paired t-test between rate at 0, rate at 1 · *L*_0_) as initial response slope is near zero. (**e**) A model with a subset of inhibitory cells stimulated (60%) can recapitulate the data in (b). Shaded region: ±1 s.d. via data bootstrap as in Fig. 4. Other than splitting inhibitory population into two subsets, network parameters are as inferred in Fig. 4 (Methods). (**f**) Population responses for transgenic expression of ReaChR in PV neurons. Unlike the viral-expression data (d), the identified inhibitory cells in these experiments show paradoxical suppression; mean firing rate is significantly suppressed (*p* < 10^−6^, paired t-test on initial slopes). (**g**) Cell counts in histological sections. Solid/dotted black lines: means across two independent human counters; upper and lower gray boundaries give results from each counter. X-axis shows variation as counters were asked to use a high or low threshold for accepting an opsin positive cell. Cells were counted across 400 *μ*m depth and are expressed (y-axis) as a percent of DAPI-positive nuclei. Transgenic expression gives about twice as many opsin expressing cells as viral expression, in addition to the differences in expression heterogeneity seen in (b-c). (**h**) Example viral (PV-Chronos) units show diversity of responses to stimulation. Some narrow spiking units (blue, rightmost two panels) show non-paradoxical initial increases, and some show paradoxical initial suppression. (**i,j**) Distribution of spike widths (i) and initial slopes (j) for experiments using transgenic PV-ReaChR (bottom). Same conventions as in Fig. 5, except here colors are as shown in panel (a). (**k,l**) Same as (i,j) for viral expression. Though inhibitory units are classified by response at high laser power, differences in spike width are visible in both datasets. Viral mean and median slopes are zero or positive (t-test for negative mean *p* > 0.05, Mann-Whitney U for negative median *p* > 0.05; 22/42 (53%) negative I slopes); transgenic mean and medians are negative (negative mean *p* < 10^−7^, negative median *p* < 10^−7^; 24/27 (89%) negative I slopes). (**m**) Summary of mean and median inhibitory cell initial slopes for these data and data from V1, somatosensory, and motor cortex.

We first used viral methods to express an opsin in PV neurons, injecting a Cre-dependent adeno-associated virus (AAV) encoding an excitatory opsin (Chronos, [36]), into a transgenic mouse line (PV-Cre [37]). In these experiments, the network neurons can be divided into three populations (Fig. 6a): (1) excitatory, (2) Chronos-expressing PV inhibitory (PV-Chronos), and (3) remaining inhibitory neurons: non-PV (e.g. somatostatin-positive, etc.) or non-Chronos-expressing PV neurons. We identified PV-Chronos cells by measuring whether cells’ firing is increased at high laser intensity (statistically significant increase at maximum laser power, as in Fig. 5; see Methods). In V1 VGAT-ChR2 experiments, this classification by response at high laser power produces qualitatively similar measures of paradoxical suppression as pharmacology-based classification, Fig. S8). Non PV-Chronos cells (without increase in firing at high laser intensity), are likely either excitatory cells, non-PV, or non-expressing cells Supporting the idea that our classification approach identifies PV-Chronos cells, the majority of classified inhibitory cells have narrow waveforms (Fig. 6i). While not all PV-positive cells are basket cells [38], the large proportion of narrowwaveform units we found suggest our inhibitory classification detects a number of fast-spiking cells.

Stimulating the PV cells produced no significant paradoxical effect on average (Fig. 6d, blue). From a theoretical point of view, this average inhibitory response is an important measure, as with the standard two-population ISN model, e.g. [10], it is the average response (averaged over inhibitory cells) that is paradoxically suppressed when a network is inhibition-stabilized and strongly coupled. A second important measure, e.g. for experimentalists wishing to identify inhibitory neurons, is how many individual inhibitory cells show paradoxical effects. Examining individual units showed that PV-Chronos units often showed no paradoxical effect: initial slope was often positive and thus non-paradoxical (e.g. Fig. 6h; median not sig. dif. from zero; summarized in Fig. 6k-l; see legend for statistical tests.).

One reason why stimulation with viral expression might produce no paradoxical effect on average is that viral expression could target only a subset of the PV cells [20]. We first examined the effect of inhibitory cell subset stimulation in a model, the rate model of Fig. 4, using network parameters inferred there. Stimulating a subset of inhibitory neurons in the model (60%) reduced the magnitude of the average paradoxical effect (Fig. 6e, analytical derivation in Methods, and Fig. S15), and described the data (Fig. 6d) well. We also considered a model with multiple subclasses of inhibitory neurons, using the connectivity structure measured in [39], and found there also that paradoxical effects were not clear when approximately half of the PV cells were stimulated (Fig. S15).

To test these models, and experimentally determine whether differences in number of cells expressing opsin could change the paradoxical effect, we used a transgenic approach to express a different excitatory opsin in PV cells (PV-Cre;ReaChR transgenic mice). In these mice, the opsin is expressed in most or all PV cells [40]. (Since we study steady-state firing rates (Methods), we do not expect differences in the onset or offset kinetics of the opsin to affect these measurements, and we verified that considering different time windows of the steady-state response does not change the results, Fig. S13.) Because we measured responses for a range of light intensities, differences in viral and transgenic mean levels of opsin expression would not affect paradoxical effects. On the other hand, differences in the number of expressing cells, or variability in opsin levels across cells, are predicted to change the paradoxical effect. Indeed, we found that, with transgenic expression, the average paradoxical effect was present (Fig. 6f). Further, the fraction of inhibitory cells that showed paradoxical responses was significantly larger in PV-ReaChR transgenic animals than in PV-Chronos animals (*cf* Fig. 6j,l), and was similar to the VGAT-ChR2 data in V1, somatosensory, and motor/premotor cortex (Fig. 6m).

Changes in the number of stimulated cells could be affected by the virus infecting only a subset of neurons [41], or by the virus yielding different levels of opsin expression in different neurons, so that only a subset are recruited strongly at any given light intensity. To examine differences in expression pattern,we counted neurons in histological sections from both the viral (PV-AAV-Chronos) and transgenic (PV-ReaChR) animals. We observed both effects: viral expression yielded opsin in about half as many cells as transgenic expression (Fig. 6g; positive-opsin cell percentage significantly different between viral and transgenic cases, both observers and both thresholds, *p* < 0.05, *χ*^2^ test), and also yielded some cells with strong, and some cells with weak, opsin expression (Fig. 6b-c). The differences in the number of opsin-expressing cells roughly matched our model predictions (60% of neurons stimulated, Fig. 6e).

Taken together, these data and our analysis support the idea that V1 (and other areas) of the mouse cortex are strongly coupled and operate as an ISN. Yet, whether an average paradoxical effect is seen depends on the number of inhibitory cells stimulated. The heterogeneous responses we saw in PV-Chronos experiments (about half of inhibitory cells paradoxical, half non-paradoxical, Fig. 6d,h,l) also may in part explain why paradoxical inhibitory suppression (and thus ISN operation) has not been more widely reported in optogenetic PV-stimulation experiments (also see Discussion).

### Paradoxical effects are also seen in deep-layer recordings

To this point all the neurophysiological data we have reported is from the upper layers (L2/3 and 4) of the cortex, recorded within 400 μm of the cortical surface. We focused on the superficial layers because the blue light we used to activate ChR2 and Chronos does not penetrate more than a few hundred microns into the tissue [42], so the inhibitory neurons that receive direct optogenetic input are those in the superficial layers. However, blue light delivered to the cortical surface to stimulate inhibitory neurons has been seen to suppress activity across cortical layers, presumably due to polysynaptic effects[43]. Because we recorded data using silicon probes that span most of the depth of the cortex, we could also assess whether light delivered to the top of the cortex also produced paradoxical suppression in deeper inhibitory neurons.

We examined units (Fig. 7) recorded ≥ 500 *μ*m from the cortical surface (see Methods). In our multi-area recordings from superficial layers (e.g. Fig. 5), we classified inhibitory cells in different brain areas based on *increases* in inhibitory response at strong light intensities, when excitatory cells are silent or largely suppressed. But in deep recordings, presumably because of light scattering and absorption, even at our highest stimulation intensities we did not always see clear mean *increases* in inhibitory firing rates (e.g. Fig. 7b). Therefore, for these deep recordings we classified neurons as inhibitory based on waveform width. As in superficial layers, we found bimodal distributions of waveform width in our deep layer data (Fig. 7a-c, insets). In our upper-layer V1 data, we compared classification of V1 units via pharmacology, waveform width, and responses at high power (Fig. S8). Based on this comparison, in these deep-layer data, we do not expect excitatory cells to be classified as inhibitory (i.e. few or no excitatory units have narrow waveforms), even though some inhibitory cells with wider waveforms will be missed with this approach (i.e. some inhibitory units have wide waveforms).

**FIG. 7:**
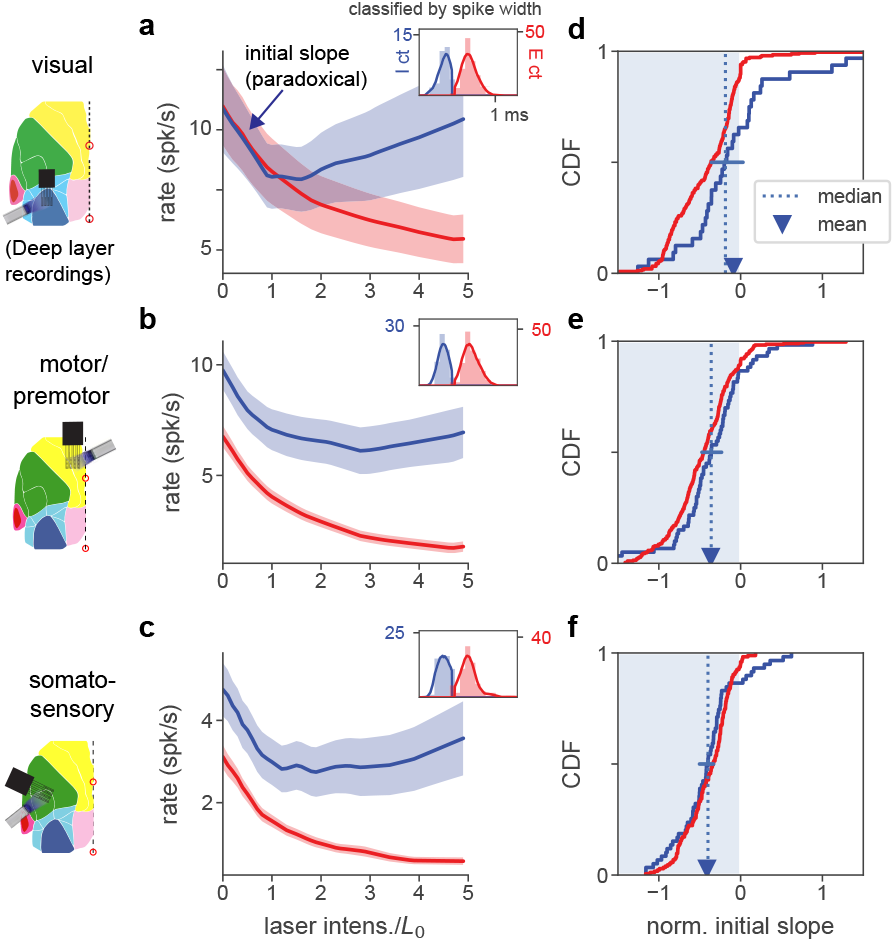
Paradoxical inhibitory suppression is also seen in deep-layer recordings. (**a**) Population average responses of deep-layer (recorded ≥ 500 *μ*m from cortical surface) units, classified as inhibitory (blue) or excitatory (red) by waveform width (inset; solid line is kernel density fit to underlying histogram, expressed in units of counts (y-axis), x-axis is spike width in ms, see Methods). Shaded area shows ±1 SEM about mean. *L*_0_ is defined for all units recorded in a single session based on responses to superficial-layer recordings (Methods). Initial slope of inhibitory average response is negative (paradoxical). (**b,c**) Same, for motor/premotor and somatosensory. (**d-f**) Initial slopes of all recorded units shown in a-c. Conventions as in Fig. 2k. As in that panel, slopes here are normalized by baseline rate so that minimum slope is −1. Means and medians of individual inhibitory neurons’ slopes are all negative, except in V1 this effect is not significant (t-test *p* < 0.01, except in V1 *p* > 0.05, blue error bar shows 95% confidence interval around median via bootstrap: upper limit V1 0.03, motor −0.33, somato −0.22). While this means there is not a clear majority of significantly suppressed units in V1, the population firing rate decrease is significant for all three areas (*p* < 0.001, Mann-Whitney U on summed population counts, baseline vs rate at *L*_0_).

Examining cells recorded from deeper layers, we found that V1, motor, and somatosensory areas all showed initial paradoxical responses (Fig. 7). However, the maximal suppression of inhibitory and excitatory cells was not as strong as in superficial layers (*cf* Figs. 2,5). (Supporting the idea that this weaker effect of stimulation is due to attenuation of blue light, paradoxical effects in deep layers were smaller in these data than when using red light to stimulate ReaChR in V1 PV neurons, Fig. S14.) In all these cases (Fig. 7a-c), the mean population inhibitory responses showed initial suppression. With the caveats discussed above (light penetration and classification of inhibitory units by waveform), these data are consistent with the idea that ISN operation is a common feature of cortical areas and layers.

### No transition out of inhibition stabilization is seen during anesthesia, at lower network firing rates

Theoretical studies have pointed out that a network can switch from non-ISN to ISN when external inputs are increased [13] If the cortical network did transition out of the ISN state at lower activity levels, this would be computationally important, as the way inputs sum can change depending on whether the network operates as an ISN or not [13; 14]. Our results above (Fig. 2,3) show that superficial layers of mouse V1 operate as an ISN at rest (i.e. without sensory stimulation), and also show the transition from ISN to non-ISN is below the level of spontaneous activity, as network activity is decreased by optogenetic stimulation of inhibitory neurons. (The transition point is where the slope of I rates vs stimulation intensity switches from being negative to positive.) However, we wished to determine whether a brain state with lower activity levels, as that produced by light anesthesia, might result in a transition to a non-ISN regime.

Under light isoflurane anesthesia, we found no evidence of a transition, and instead found that paradoxical inhibitory suppression was maintained (Fig. 8a). At 0.25% isoflurane (a low level, as surgical levels are often 1.0% and above), spontaneous firing rates of excitatory neurons are reduced (Fig. 8c; mean 6.4 spk/s reduced to 3.5 spk/s, *p* < 0.02, Wilcoxon signed-rank test; inhibitory neurons’ rates show a negative trend but are not statistically different, 14.5 spk/s awake to 11.6 spk/s anesth., *p* < 0.10, Wilcoxon). Thus, the network changes induced by anesthesia do not cause the network to transition out of the ISN state. At this low level of anesthesia, we did not observe prominent up and down state slow oscillations. In one experiment, we used a higher level of anesthesia (0.5% isoflurane, Fig. 8b), yielding even lower firing rates but still preserving the paradoxical effect. Further confirming the robustness of coupling to changes with anesthesia, the distribution of response slopes is roughly unchanged (Fig. 8c), suggesting the network is far from a transition into a non-ISN state. Anesthesia thus preserves the paradoxical inhibitory response, leaving the network still an ISN.

**FIG. 8:**
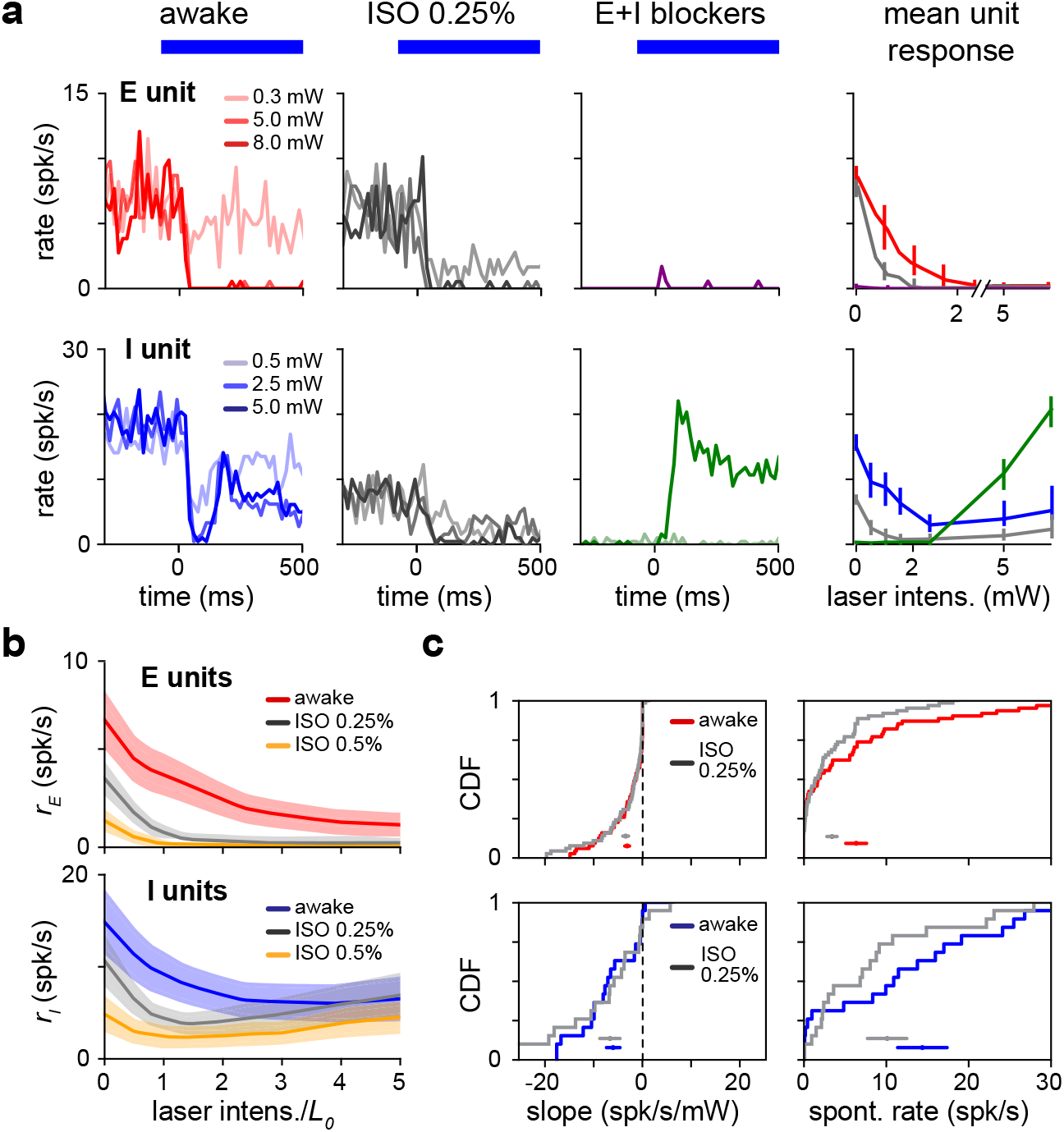
Paradoxical response is preserved with lower network activity due to anesthesia. (**a**) Single excitatory (top) and inhibitory (bottom) unit response in the awake state (no anesthesia, no synaptic blockers), under anesthesia with no blockers (isoflurane, 0.25%) and with synaptic blockers (CNQX, APV, bicuculline; Methods) plus anesthesia. (**b**) Population average response with (gray) and without (blue;red) anesthesia. Spontaneous firing rates are reduced for both the E (top) and I (bottom) populations below the awake firing rates, yet inhibitory paradoxical suppression is preserved with anesthesia (lower panel: gray line shows negative initial slope). In these experiments, inhibitory responses are weaker at high powers, but this does not affect paradoxical suppression. (**c**) Distribution of initial slope and spontaneous rate before and after anesthesia. Initial slope is largely preserved while excitatory and inhibitory spontaneous rates are suppressed. Colored horizontal bars within each panel show mean±SEM for each distribution. Initial slopes are plotted here non-normalized (units spk/s/mW) to show the slopes are quantitatively similar across firing rate changes; normalized slopes are shown in Fig. S16, and their means and medians remain negative across anesthesia state.

## III. DISCUSSION

These data show a signature of inhibitory stabilization, the paradoxical suppression of inhibitory cells to optogenetic excitation, in several different areas of mouse cortex, V1, S1, and motor cortex, during spontaneous activity in the absence of sensory stimuli. To study whether these cortical networks operate in the ISN regime, and are thus strongly coupled, we used a mouse line where all inhibitory cells express opsin. We found clear paradoxical effects in response to inhibitory stimulation. We also found, as predicted by ISN models, that strong inhibitory stimulation causes the network to transition into a non-ISN state. In this non-ISN state, inhibitory responses are non-paradoxical and increase their firing when stimulated, and excitatory neurons are suppressed. This transition from ISN to non-ISN behavior we observed is a prominent prediction of ISN models. We see paradoxical suppression of the mean inhibitory firing response in both superficial and deep layers of the three areas, and we find that a clear majority of inhibitory units (except in one of these six measurements, deep layers of V1) show paradoxical suppression when all inhibitory cells are stimulated. We use several ISN rate models to fit data from V1, aided by pharmacological manipulations that yield additional informative data about the network. We find that despite having fewer parameters than data degrees of freedom, the ISN model describes the data well. During light anesthesia, mean V1 network firing rates decrease, but we still observe ISN paradoxical suppression. Finally, we explored the effect of stimulating a subclass of inhibitory neurons with strong network effects: the PV neurons. We find that stimulating PV cells also produces paradoxical effects. We compared viral and transgenic expression strategies and found that viral expression did not produce average paradoxical effects. This viral/transgenic difference is consistent with theoretical observations that changing the fraction of stimulated inhibitory cells changes the strength of the paradoxical effect, and is supported by histological data. Subclass-stimulation paradoxical effects do not imply inhibition stabilization ([19; 44] and see Mathematical Methods), and so our PV data alone cannot determine whether the network is an ISN. However, our all-inhibitory stimulation experiments establish the network operates in a strongly-coupled ISN regime.

Our data and theoretical results provide the strongest evidence currently available to rule out non-ISN interpretations. For example, an alternative to the ISN explanation for our data might be a model with multiple inhibitory populations and weak recurrent coupling, perhaps driven by strong external input such as thalamic or corticocortical input. In principle, such an alternative model could show paradoxical-like inhibitory responses, where stimulation of one inhibitory subclass (e.g. PV cells) inhibits a second set of inhibitory cells whose firing rate decreases (e.g. somatostatin or SOM cells; the same explanation could hold with the PV and SOM classes swapped). Our results supply converging observations that rule out such an explanation — that is, they rule out a non-ISN, feedforward inhibitory-onto-inhibitory (disinhibitory), explanation. Three prominent features of our data supporting a strongly-coupled recurrent origin for our data are: (1) that inhibitory response dynamics match ISN predictions, (2) that an ISN model describes the data well and makes predictions verified in the data, and (3) many inhibitory cells show paradoxical effects at low stimulation intensity but increase their firing rate at higher stimulation intensity. In a feedforward or disinhibitory model, there would be little reason to predict inhibitory neurons to change from decreasing to increasing rate as stimulation intensity increases, but ISN models are predicted to transition into a non-ISN state that shows these effects. Our results thus support the idea that the cortex operates as an ISN even at rest.

In the last decade, multiple experiments have performed optogenetic stimulation of cortical interneurons, but these studies have yielded mixed results on inhibition stabilization. In layer 2/3 of mouse V1, optogenetic excitation of PV cells with viral expression was shown to generate non-paradoxical modulation of inhibition [17]. In mouse auditory cortex (A1), in contrast, Kato et al. [15] found that suppression of inhibitory cells with viral ArchT produced paradoxical effects in intracellular currents. However, Moore et al. [16] reported, also in A1, that stimulation of PV cells using viral transfection produced paradoxical effects in some PV cells and not others, and they reported evidence for disinhibitory, non-ISN, mechanisms via L4 inputs into L2/3. And a recent study in somatosensory and motor cortex observed paradoxical suppression for PV subclass stimulation in some layers and not others [44]. Finally, many experiments have used excitation of inhibitory cells (e.g. [34]) to suppress excitatory activity, without reporting paradoxical inhibitory effects, though a recent survey of such methods does report paradoxical effects [43] in somatosensory and motor cortex. Our results, combining theoretical analyses with a range of experimental conditions, suggest some explanations why paradoxical inhibitory suppression has not been previously widely reported.

First, it appears a significant fraction of inhibitory cells must be stimulated to produce large paradoxical changes in firing rate [20]. Stimulating PV cells with viral methods (which can yield subsets of cells with weaker or no expression; see also [18; 20] for a similar explanation for non-paradoxical effects with viral expression), produced in our hands mixed effects on single cells and weaker mean paradoxical effects (Fig. 6). Further, stimulating subclasses of inhibitory cells even with transgenic expression can produce different inhibitory responses compared to stimulating all inhibitory neurons [44]. Second, especially in experiments using extracellular recording, it can be difficult to definitively identify inhibitory cells. Some inhibitory cells’ waveforms are broad (Fig. 2), and in an ISN, excitatory and inhibitory cells’ firing rate changes are often similar (inhibitory neurons may only show increased activity for a few milliseconds before paradoxical suppression begins). Third, for strong enough drive to inhibitory neurons, ISNs transition to a non-ISN state. In this non-ISN state with low excitatory activity, tracking of excitatory fluctuations is not needed to stabilize the network, and some past experiments may have used stimulation large enough for parts of the network to enter into this second phase of activity, hiding the signature of inhibition stabilization.

In an ISN, strong bidirectional E-I synaptic coupling allows inhibitory cells to stabilize an unstable excitatory network, by tracking and responding to changes in local excitation. While paradoxical effects in a single inhibitory subclass do not imply overall network stability (Fig. 6e), PV neurons may well be the principal providers of stabilizing input. PV neurons receive input from many diverse local excitatory cells [45], show responses that reflect an average of local excitatory responses [46; 47], cause network instability when strongly suppressed (an effect which does not result from SOM suppression) [48], and target virtually all nearby excitatory cells with strong peri-somatic synapses [49; 50]. In principle, other cell types of the diverse inhibitory classes in the cortex [25] could contribute to stabilization, though evidence suggests specific circuit effects (e.g. gain control) may be controlled by other classes of inhibitory cells (e.g. VIP, SOM).

Several influential models of cortical function include inhibition-stabilized parameter regimes, and our data provide new constraints on such models. The ‘balanced network’ model predicts that both excitatory and inhibitory inputs to single neurons should be very large, but approximately cancel each other, leading to a balance between excitation and inhibition [5; 7]. This scenario accounts in a parsimonious way for multiple ubiquitous properties of activity in cortex, such as the highly irregular nature of neuronal firing, and the broad distributions of firing rates across neurons. Another influential model of cortical operation, the SSN model, can be described [13] as a “loosely balanced” model with moderate recurrent coupling [33]; increasing its coupling strength leads to a balanced network model. The network parameters that best fit our data produce moderately large (compared to threshold) excitatory and inhibitory inputs of approximately similar magnitudes (see Fig. 4), a finding consistent with loose balance.

The SSN makes a specific prediction about the ISN regime that our data addresses. The SSN predicts that cortical networks are inhibition-stabilized for strong inputs, but not an ISN for sufficiently weak inputs. Thus, the SSN predicts a transition between two operating regimes as network activity increases. Finding this transition point is important for understanding computation, as the two regimes show different (supralinear or sublinear) modes of input summation. Prior ISN studies [12] have left open whether this transition occurs only with sensory stimulation. A study of responses to two combined sensory stimuli [51] found a linear-to-sublinear transition for increasing strength (contrast) of visual stimuli, but did not resolve whether the non-ISN to ISN transition was above or below the level of spontaneous activity. Our data do resolve this, showing several cortical networks are in the ISN state even without sensory stimulation. But a second question our data address is whether during spontaneous activity the network is on the edge of this transition point, or well into the ISN regime. Our work gives two pieces of evidence that without sensory activity the network is far above a transition into a non-ISN state: first, excitatory rates must be substantially suppressed before inhibitory responses switch from paradoxical suppression to firing rate increases (e.g. Fig. 2e), and second, under light anesthesia, we find network activity is lowered but ISN behavior is preserved. Therefore, our data provides support for the idea that cortical areas generally operate above this transition point, and that the weakly-coupled, non-ISN regime predicted by the SSN is not a regime mouse cortex commonly enters during normal processing in awake behavior.

Together, the results reported here suggest that inhibition stabilization is a ubiquitous property of cortical networks. This data is consistent with a network that is in the strong coupling regime, but is not extremely strongly coupled as some balanced network models would predict [5; 7]. It is tempting to speculate that, while strong coupling allows the network to perform non-trivial computations on its inputs, increasing coupling further might not be optimal for several reasons. First, in the strong coupling limit, only linear computations are available to a balanced network (unless synaptic non-linearities are included, [52]), while networks with moderate coupling can combine inputs in a non-linear fashion [13]. Second, maintaining strong connections might be metabolically expensive, so that a moderately coupled network might represent an ideal compromise for cortical computations.

## ACKNOWLEDGMENTS

Supported by the NIMH Intramural Research Program and by NIH BRAIN U01 NS108683 (to M.H. and N.B.). We thank K. Miller and J. Reynolds for discussions and the NIMH Instrumentation Core for technical support. We also thank L. Glickfeld, S. Lee and K. Miller for comments on the manuscript. This work used the computational resources of the NIH HPC Biowulf cluster (http://hpc.nih.gov).

## IV. METHODS

### A. Experimental methods

#### 1. Animals

All procedures were conducted in accordance with the guidelines of the National Institutes of Health. Seven VGAT-ChR2, three PV-Cre, and two PV-Cre;ReaChR mice were used (JAX stock #s 014548, 008069, and 024846; 2 females and 10 males; singly housed on a reverse light/dark cycle).

#### 2. Cranial window implants

Mice were implanted with a titanium headpost and a transparent window (optical glass, 0.8 mm thickness, 3 or 5 mm diameter) in the left cerebral hemisphere. The windows provided access to the primary visual (V1) and somatosensory cortex, or motor cortices, for imaging and silicon neural probe recordings with optogenetics. Mice were given dexamethasone (3.2 mg/kg, i.p.), 2 hours before surgery. Animals were anesthetized during surgery with isoflurane (1.0%-4% in 100% O_2_). Using aseptic technique, a headpost was affixed using C&B Metabond (Parkell), and a 3 or 5 mm diameter craniotomy made.

#### 3. Hemodynamic intrinsic imaging

To determine the location of V1, we delivered small visual stimuli to animals at different retinotopic positions and measured changes in the absorption of 530 nm light resulting from cortical hemodynamic responses [53]. We evenly illuminated the brain with a 530 nm fiber coupled LED (M530F2, Thorlabs) passed through a 532 nm-center bandpass filter (Thorlabs). Images were collected on a stereo microscope (Discovery V8, Zeiss) through a green long-pass emission filter using a 1x objective (PlanApoS 1.0x, Zeiss) onto a Retiga R3 camera (Q Imaging, captured at 2 Hz with 4×4 binning). For retinotopic mapping we presented upward-drifting square wave gratings (2 Hz, 0.1 cycles/degree) masked with a circular window (10°diameter) for 5 s with 10 s of mean luminance preceding each trial. Stimuli were presented in random order at four positions in the right monocular field of view. The response to a stimulus was calculated as the fractional change in intensity between the average of the 10 frames immediately preceding the stimulus (as baseline) and a 6-10 frame window 1 s after stimulus onset (as response) to match the timecourse of the hemodynamic response [54; 55].

#### 4. Viral injections

We used viral injections to express Chronos [36] in PV-Cre [37] animals. Mice were anesthetized (isoflurane 1-1.5%) and the cranial window implant removed. We used a stereotaxic injection system (QSI, Stoelting Inc.) to deliver 500 nL of a 10:1 mixture of AAV1-hSyn-FLEX-Chronos-GFP (UNC Vector Core Stock) and 100 *μ*M sulforhodamine (SR101, Invitrogen, for visualization) at a depth of 200-400 μm. We made 3 to 4 injections spaced 0.5-1.0 mm apart to cover the visual cortex. After the injections a new cranial window was affixed. Viral expression was monitored over the course of days by imaging GFP fluorescence, and allowed to develop for greater than 4 weeks before electrophysiological recordings.

#### 5. Electrophysiological recording with optogenetic activation of inhibitory neurons

For recording experiments, we first affixed a 3D printed ring to the cranial window to retain fluid. With the ring in place, we removed the cranial window and flushed the craniotomy site with sterile normal saline to remove debris. Kwik-Sil silicone adhesive (World Precision Instruments) was used to seal the craniotomy between recording days. Using a stereomicroscope on an articulating arm we positioned the end of an optical fiber (600 μm diameter, Doric Lenses) fitted with a light-tight coupler to an optical cannula (400 μm diameter, Thorlabs CFMLC14L02) over our target cortical area, with a slight 10-30° angle from vertical to provide space for the electrodes. We used fiber coupled LED light sources (M470F3 (Chronos and ChR2) or M625F2 (ReaChR), Thorlabs) to deliver illumination with peak at 470 nm or 625 nm to the brain. We calibrated total intensity at the entrance of the cannula using a power meter and photodiode (meter model 1918-R, photodiode model 918D-SL-OD3R, Newport Corp.). The cannula distance from the dura was adjusted to provide a light spot with a full width at half maximum intensity of 0.8-1.2 mm (as measured with a small digital macro documentation microscope, Opti-TekScope); thus, our reported spot irradiance in mW/mm^2^ is within a factor of two of the reported power (mW). We targeted a multisite silicon probe electrode (NeuroNexus; 32-site model 4×8-100-200-177; 4 shank, 8 sites/shank; sites were electrochemically coated with PEDOT:PSS [poly(3,4-ethylenedioxythiophene): poly(styrenesulfonate)] [56]) to the center of the LED spot using a micromanipulator (MPC-200, Sutter Instruments). With the fiber optic cannula and electrode in place, we removed the saline buffer using a sterile absorbent triangle (Electron Microscopy Sciences, Inc.) and allowed the dura to dry for 5 minutes. After insertion, we waited 30-60 minutes without moving the probes to reduce slow drift and provide more stable recordings. We isolated single and multiunit threshold crossings (3 times RMS noise) by amplifying the site signals filtered between 750 Hz and 7.5 Khz (Cerebus, Blackrock microsystems). During recordings animals were awake and passively viewing a gray screen. To keep animals awake and alert, animals were water-scheduled [57], and a 1 *μ*l water reward was randomly provided on 5% of the stimulus trials; we verified animals were licking in response to rewards during the experiments. Superficial units were those recorded from a site within 400 *μ*m of the cortical surface, identified by monitoring spike and local field potential activity (LFP) on sites at different depths; we typically observed desynchronized cortical LFP activity on any site below the surface and found the first substantial unit activity 100 *μ*m below the surface. Deep units were those recorded 500-800 *μ*m below the surface. Optogenetic stimuli were square light pulses with 2 ms linear ramps at start and end to reduce recording artifacts. Pulses were on for 600 ms and off for 1000 ms at a range of power levels (0.3-10 mW), presented in random order, with 100 repetitions per power level. For population plots, we normalized the light intensity in each experiment to control for fluctuations across experiments due to e.g. changes in dural thickness or tissue light absorption. We found a minimum value of the inhibitory responses (*L*_0_, with one value for each experiment, used for each unit recorded in that experiment) by fitting a 2-segment piecewise-linear function to the average inhibitory response (Fig. S4). To avoid biases due to light attenuation at deep sites, *L*_0_ was computed for each experiment on the superficial sites only, and this value was used for the deep site data (Fig. 7). Population firing rate std. dev. and SEM (Fig. 3) were calculated on the sum of all unit counts in a given time bin.

#### 6. Pharmacological blocking of excitatory and inhibitory synapses

To classify cells using pharmacological weakening of cortical synapses, we divided the experiment into three phases, each 30-45 minutes in duration: first no blockers, then excitatory blockers, then excitatory plus inhibitory blockers. For each section, we delivered the same optogenetic stimulation protocol (above). To apply the pharmacological agents, we made a hole near the recording electrodes in the agarose on top of the brain, removed the normal saline covering the agarose and hole, and replaced it with a solution containing the agent in saline. After 15 minutes, the initial application of blocking solution was removed via aspiration and refreshed. We waited a total of 20 mins for the pharmacological agent(s) to take effect before recording. Excitatory synaptic blockers (2 mM CNQX, 6 mM APV) affected AMPA, kainate, and NMDA synapses. Inhibitory blockers (1 mM bicuculline) affected GABA-Asynapses. Source: Sigma-Aldrich (#C239, #A8054, and #14343).

#### 7. Histology and electrophysiology probe tracking

To determine the location and depth of recording electrodes we coated the shanks with 1,1’-Dioctadecyl-3,3,3’,3’-tetramethylindocarbocyanine perchlorate (DiI) (Thermo Fisher #D282, 50 mg/mL solution in ethanol), prior to insertion into the brain. We dipped the electrode tips 5 times into the DiI solution and allowed the coating to dry for 30 seconds between immersions. Fluorescent electrode tracks were then visualized in coronal sections of fixed brain tissues. Mice were anesthetized with isoflurane and injected intraperitoneally with pentobarbital sodium (150 mg/kg). They were then perfused transcardially with cold (4C) PBS followed by cold 4% paraformaldehyde. Brains were extracted and fixed in 4% paraformaldehyde for 6-12 hours and then cryoprotected in a 30% (w/v) sucrose solution in PBS until they sank. Brains were sectioned at 50 μm on a freezing microtome (Leica), mounted on glass slides, and coverslipped with mounting media containing DAPI (Fluoromount-G with DAPI, Electron Microscopy Sciences). Slides were imaged using an Olympus slide-scanner (Olympus BX61VS, Japan).

#### 8. Counting opsin-expressing cells

To quantify opsin expression differences between viral (PV-Cre,AAV-Chronos-YFP) and transgenic (PV-Cre::ReaChR-mCitrine) animals, we used a confocal microscope (Zeiss LSM780, 20x objective) to image 50 *μ*m thick coronal brain tissue sections. We imaged DAPI and either YFP or mCitrine fluorescence from layer 2/3 of the visual cortex. For each animal, we chose four 100 *μ*m x 100 *μ*m areas in layer 2/3, and constructed 3D stacks by aligning images across adjacent sections. 3D volumes from 3 viral and 3 transgenic animals were anonymized and manually counted by two independent observers. Observers were instructed to count cells once with a high threshold for accepting a cell as positive, and once with a low threshold for accepting a cell. Percentages of opsin-expressing cells were calculated as the average number of fluorescently labeled cells divided by the number of DAPI-labeled nuclei.

#### 9. Anesthesia during electrophysiology

To test the effects of lowering overall activity on the stability of the cortical network we fitted animals with an isoflurane inhalation mask system (model V-1, VetEquip Inc.) and provided anesthesia (0.25-0.5%isoflurane in 100% O_2_) to put the animal into a lightly anesthetized but awake state during recordings. At the lower 0.25% isoflurane concentration, recordings in V1 displayed no synchronized oscillatory Up/Down state activities and the animals eyes were open throughout. The anesthesia experiments were divided into four sections: awake animal, anesthesia with no blockers, anesthesia with excitatory blockers, anesthesia with inhibitory blockers. Anesthesia was delivered for 20 minutes before starting recording.

#### 10. Spike sorting

Spike waveforms were sorted after the experiment using OfflineSorter (Plexon, Inc.). Single units were identified as waveform clusters that showed clear and stable separation from noise and other clusters, uni-modal width distributions, and inter-spike interval histograms consistent with cortical neuron absolute and relative refractory periods. Multiunits were clusters that were distinct from noise but did not meet one or more of those criteria, and thus these multiunits likely group together a small number of single neurons. Signal-to-noise ratio (SNR) [58; 59] of single unit waveforms (median ±std): in visual cortex, 3.6±1.7 (N=166), see Fig. S5; in other datasets: motor, 3.7 ±1.2 (N=103); somatosensory, 3.7 ±1.7; in V1 PV data (Fig. 6) 2.91 ±1.3. Only single-units were analyzed; multiunits were discarded. We also repeated our single-unit analyses using both more- and less-stringent criteria to qualify a unit as a single unit, and found no qualitative differences in the results (Supplementary Fig. 4). Supporting the idea that single units did not group together multiple units, we found no significant correlation between SNR and baseline firing rate in any of the four datasets (all *p* > 0.2 by linear regression).

### B. Mathematical methods

#### 1. Model equations

To quantitatively analyze the network response, we used a two population rate model [60], in which the average firing rates of excitatory (E) and inhibitory (I) populations (*r_E_* and *r_I_*, respectively) evolve according to

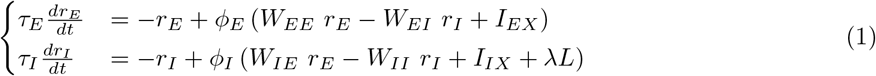

where *ϕ_A_*, *I_AX_* and *τ_X_* are the static transfer function (f-I curve), external input and time constant of population *A* (=E,I), respectively, while *W_AB_* is the strength of connections from population *B* to *A*. The optogenetic stimulation is described by the parameters λ and *L* which represent the efficacy and the intensity of the stimulation light. Results shown in the main text (Fig. 4) have been obtained with a rectified-linear transfer function

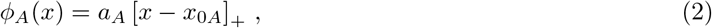

which is zero for input *x* smaller than the threshold *x*_0*A*_, and increases linearly, with a gain *a_A_*, otherwise. This transfer function is the simplest one that can describe the data, which shows an approximately piecewise linear dependence of firing rates on stimulation intensity (see Fig. 4). We also fit the data using a different transfer function

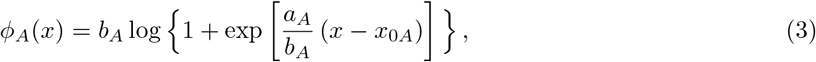

that smoothes the threshold non-linearity of the rectified linear function, using an additional parameter *b_A_* that controls the width of the exponential region around threshold. This function reduces to the rectified-linear transfer function when *b_A_* → 0. We find that the nonlinear transfer function provides a minor improvement in describing the data and does not significantly affect the values of the inferred parameters (Fig. S12).

In the experiments, recordings of the response are done in three separate phases: (1) A ‘normal’ phase in which recurrent interactions are intact; (2) A phase with E synaptic blockers; and (3) A phase with both E and I synaptic blockers. The addition of blockers in the model is described with two parameters, *ϵ_E,I_* ∈ [0,1], representing the decrease in strength of excitatory and inhibitory synapses. After the addition of excitatory blockers, connectivity is modified as

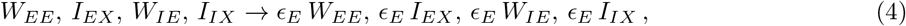

and with inhibitory blockers, connectivity is modified as

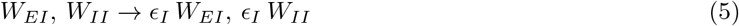

Combining Eqs.(1), (4) and (5) with the transfer function of Eq. (2), the model uses 15 parameters (2 time constants, 2 thresholds, 2 gains, 4 connectivity strengths, 2 external inputs, 1 stimulation efficacy, 2 blocker efficacies) to describe simultaneously the three phases of the experiment.

To compare the model response with the data, we find the equilibrium solution of Eq. (1), shown in Eq. (6). Since *τ_E_* and *τ_I_* do not affect the equilibrium solution, the number of relevant independent variables in the model reduces to 13. Moreover, the values of *a_E_* and *a_I_* can be reabsorbed in the definitions of *W, I* and λ. It follows that all the parameters can be inferred up to a proportionality constant and, without loss of generality, we can fix *a_E_* = *a_I_* = 1. This reduces the number of independent parameters to 11.

Note that this number of parameters is smaller than the number of parameters needed for piecewise linear fits of the data (average firing rates vs light intensity for both E and I neurons). Such piecewise linear fits of the data need 15 parameters: 3 phases times 5 parameters per phase (3 for inhibitory neurons - 2 per linear region, minus one for the continuity constraint; and 2 for excitatoryneurons, for the single linear region at low intensities). Thus, the model does not have enough parameters to fit successfully the data, unless the model structure accurately describes the cortical responses. Indeed, Fig. 4 shows that the model gives a good description of the data. The additional 4 degrees of freedom present in the data lead to four parameter free model predictions described in the main text.

The fixed point solutions of Eq. (1) (without blockers) are

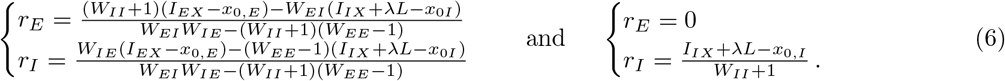

For every value of the laser intensity, the excitatory and inhibitory nullclines are defined as the functions *r_I_* = *r_I_*(*r_E_*) which solve the first and the second line of Eq. (6), respectively. Every intersection between the two nullclines is a solution of Eq. (6) and gives a stationary state of the network dynamics.

Eq. (6) has two dimensions that can be inferred only up to a multiplicative constant:

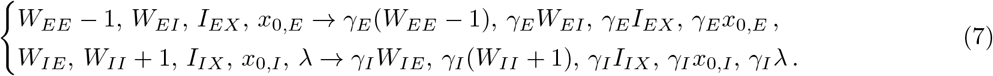

Because of these invariances, using data from a single experimental phase, only ratios between parameters can be inferred. This is no longer true when the three phases are considered; in this case all the 11 relevant model parameters can be found.

#### 2. Parameter inference

Given the dataset of excitatory and inhibitory responses, the best set of parameters describing the data is found via global optimization as follows:

1. Select random initial parameters in the interval [0,10].
2. Find the optimal set of parameters through a least-squares optimization (python function “curve_fit”, variables are constrained to be positive and, for *ϵ_E,I_*, in the interval [0,1]) of the difference between observed rates and model predictions given by Eq. (6).
3. Repeate the procedure 10^4^ times, and select the solution with minimal error as the optimal parameter set describing the data.

The optimization procedure is applied simultaneously on data from all phases and recording sessions in V1. For the first phase (awake), data from all recording sessions from VGAT-ChR2 animals in V1 (9 days, 4 animals) were pooled together for fitting. For the second and third phases (E blockers, E+I blockers), in order to describe the variability from day to day of the efficacy of the blockers, we use a different set of *ϵ_E,I_* for each recording session (6 days, 2 animals). For these two phases, 3 out of the 9 recording sessions are excluded from the analysis because blockers were added after anesthetizing the animals. The optimal parameters found by this approach are:

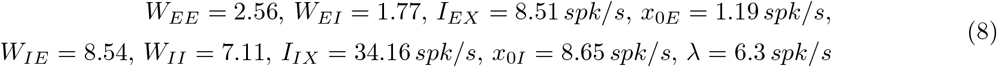

These parameters are used in Eq. (6) to generate the best model description of the data in the first phase of the experiment (Fig. 4A, first column, black line). For the second and third phases, predictions also depend on the efficacy of synaptic blockers, which are different in the different recording sessions (see Fig. S11). Data and model predictions for all sessions are shown in Fig. S11; in the main text, we showed averages of the two computed across sessions for each phase (Fig. 4A, second and third columns, black line).

We use a bootstrap approach, combined with the global optimization described above, to estimate the precision with which model parameters can be inferred from the data. The optimal (minimum-error) estimates obtained from 10^4^ random resamplings (random with replacement) are shown in Fig. S11. Despite the non-uniqueness of the solutions, the bootstrap shows (dashed lines in Fig. 4: 1 s.d. via boostrap) that model rates are clustered around the data in all three phases of the experiment.

#### 3. Stability of the solution

To analyze the stability of the system, we compute the eigenvalues of the dynamical matrix defined by Eq.(1). For the linear transfer function we find

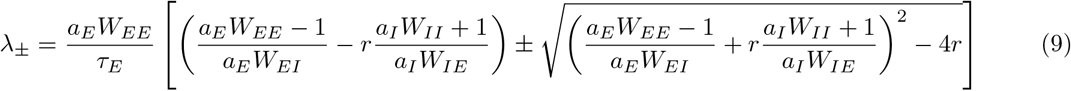

with 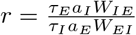 This ratio cannot be inferred from the data since *τ_E,I_* do not appear in the static solution.

We thus determined *τ_E_* and *τ_I_* (shown in Fig. 4e) from the response dynamics (Fig. 4d).

#### 4. With multiple inhibitory subclasses, ISN operation need not imply paradoxical suppression

To understand how the network response to PV stimulations depends on its structure, here we analyze a three population model (Fig. 6a) with one excitatory and two inhibitory populations, referred hereafter as *E* (pyramidal cells), *P* (ChRonos-expressing PV cells), and *I* (non-expressing PV, SOM, VIP…). The network dynamics is described by

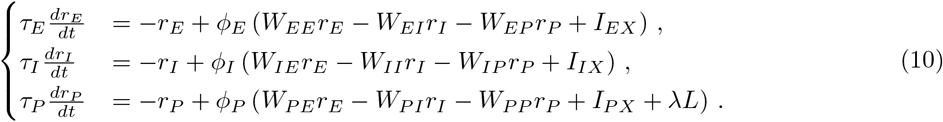

The model predictions in Fig. 6 have been obtained from Eq. 10 using

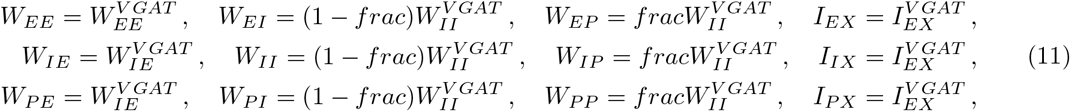

where *f_rac_* ∈ (0, 1) represents the fraction of ChRonos-expressing PV cells, 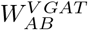 and 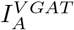 represent the parameters of Eq. (8), obtained using the V1 data from the VGAT-ChR2 mouse line.

Using the rectified linear transfer function of Eq. (2), and assuming *a_A_* = 1 to simplify expressions, the equilibrium response to stimulation of the *P* population is given by

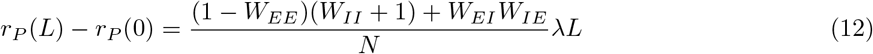

with

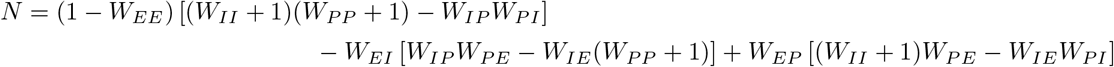

Eq. (12) shows that, both for *W_EE_* < 1 and *W_EE_* > 1, depending on the network connectivity, the ratio on the r.h.s. can be either positive or negative. It follows that, in a network with two inhibitory populations, an unstable excitatory subnetwork does not imply paradoxical suppression when one of the populations is stimulated. This can be seen more easily in a simplified model in which connectivity depends only on the identity of the presynaptic unit. In particular, using *W_EE_* = *W_IE_* = *W_PE_* = *W*, *W_EI_* = *W_II_* = *W_PI_* = *k_I_W*, and *W_EP_* = *W_IP_* = *W_PP_* = *k_P_W*, Eq. (12) becomes

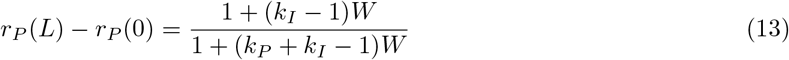

In the large *W* limit, in which the excitatory population is unstable without inhibition, the network shows the paradoxical effect only if *k_I_* < 1 and *k_P_* > 1 – *k_I_*. (Note that, for the network to be stable in this condition, the matrix of the coefficients of Eq. (10), linearized around the fixed point, must have eigenvalues with a negative real part, a requirement which also involves the magnitude of *τ_E_*, *τ_I_*, and *τ_P_*.)

## SUPPLEMENTARY FIGURES

**FIG. S1:**
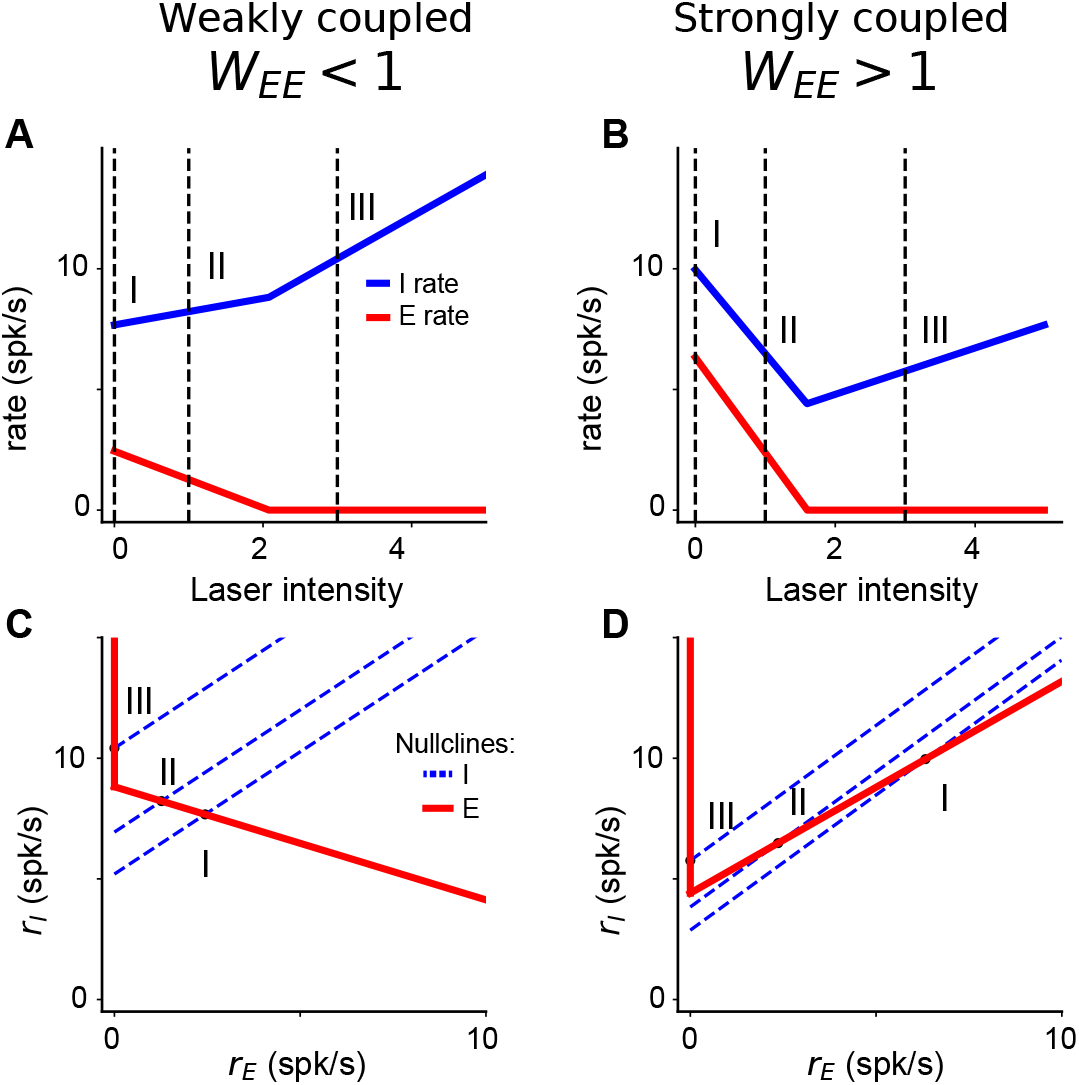
*Assoc. with Fig. 1*. Nullclines characterizing the network response at different levels of inhibitory drive. (**A-B**) Firing rate of E (red) and I (blue) populations computed from Eq. (1) as a function of laser intensity when the excitatory sub-network is stable (A, *W_EE_* < 1; weak coupling) or unstable (B, *W_EE_* > 1, strong coupling). As described in main text, inhibitory cells increase firing rate at high laser intensities in both networks. Differences are seen at lower laser intensities, where the inhibition stabilized network (*W_EE_* > 1) responds paradoxically. (**C-D**) Differences in inhibitory responses are seen in the corresponding nullclines (defined as the stationary solutions of the two lines of Eq. (1)) For *W_EE_* < 1 (C), at low laser intensity (I), the excitatory (red) and the inhibitory (blue) nullclines intersect at a nonzero value of *r_E_* (which represents the stationary point of the network). Increasing the laser intensity shifts the inhibitory nullcline to position (II) and produces a new intersection point with larger *r_I_* and lower *r_E_*. At large laser intensities (III), the two nullclines intersect at *r_E_* equal to zero and the corresponding *r_I_* value increases with the laser intensity. For *W_EE_* > 1 (D), when the laser intensity increases from (I) to (II), both the stationary value of *r_I_* and *r_E_* decrease. This happens because the excitatory nullcline is an increasing function of *r_E_*, a property which appears when that the excitatory population is unstable [10]. At large laser intensity (III), the excitatory population is silent and stable, both the excitatory and inhibitory nullclines behave as in the *W_EE_* < 1 case, and increasing the laser intensity increases the inhibitory rate.

**FIG. S2:**
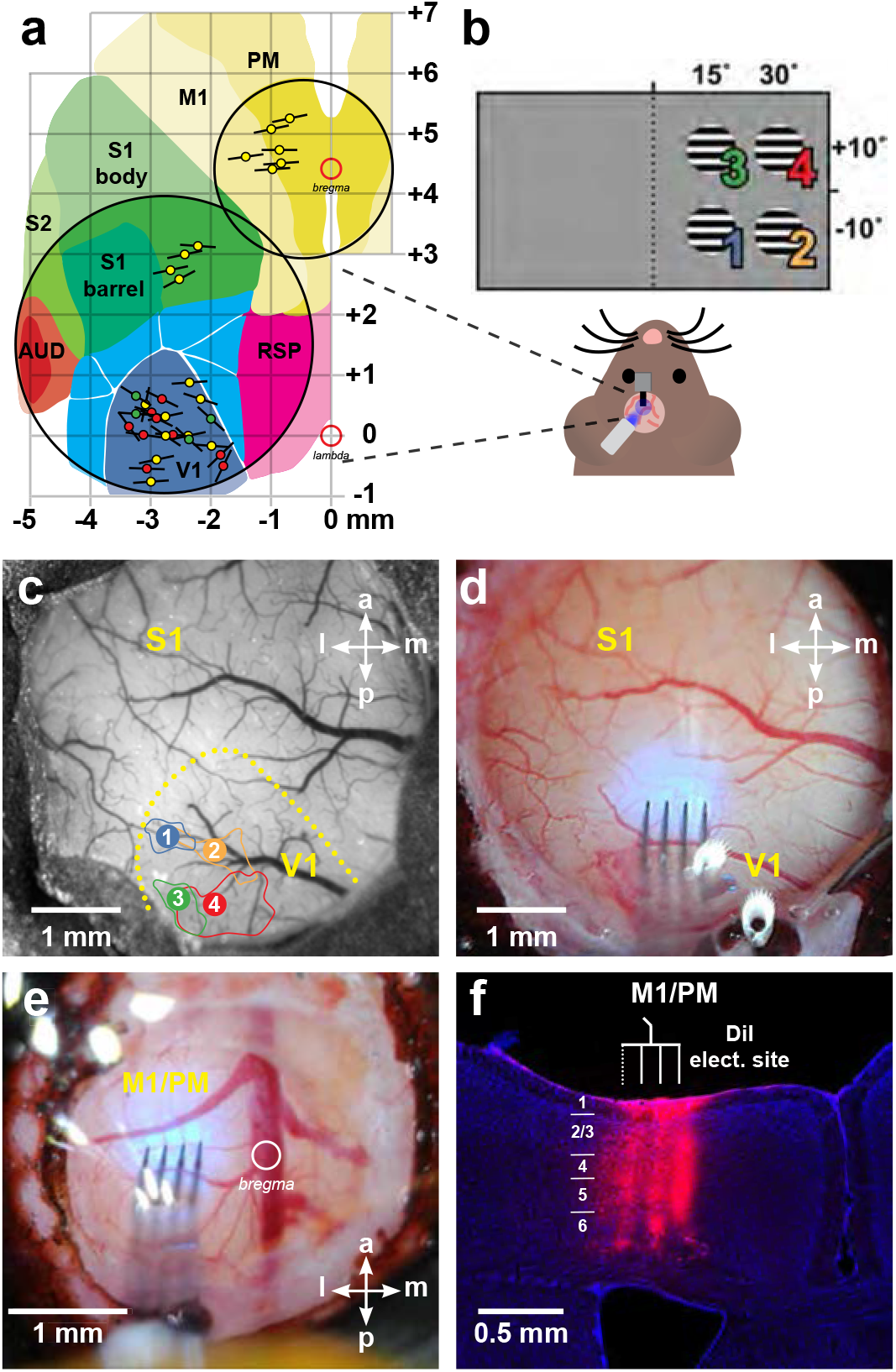
*Assoc. with Fig. 1*. Locations of recording sites. (**A**) Schematic of all physiology probe placements in the visual, somatosensory, and motor cortices (black lines give approximate extent of shanks on the 4-shank probes; colored circles signify experiment types: VGAT-ChR2: yellow; viral PV-Cre;AAV-FLEX-ChRonos: red; transgenic PV-Cre;ReaChR: green). V1: primary visual (Allen notation: VISp); RSP: retrosplenial; AUD: auditory areas; S1: primary somatosensory (SSp); S2: secondary somatosensory (SSs); M1: primary motor (MOp), PM: premotor (MOs). Coordinates are in millimeters relative to lambda and bregma (open red circles). Area boundaries drawn from Allen Mouse Brain Connectivity Atlas [61]. (**B**) Schematic of hemodynamic intrinsic-signal optical imaging (IOI) for retinotopic mapping of V1 (VISp). Hemodynamic responses were measured to upward-drifting square wave gratings (2 Hz, 0.1 cycles/°) masked with a circular boundary (10° diameter) at four locations in the right monocular field of view. (**C**) Picture of a large-diameter (5 mm) cranial window allowing access to the visual and somatosensory cortices. Yellow outline: estimated edge of V1 via retinotopic responses; colored numbers: centroid of IOI response corresponding to the stimulus positions in panel B; solid lines of the same colors: areas yielding responses within 50% of peak response. (**D**) Example physiology probe placement and blue LED overlap for optogenetic stimulation in V1. (**E**) Picture of a smaller-diameter (3 mm) window implant allowing access to the motor cortex. (**F**) Histological section showing location of electrode tracks in M1/PM. Section at 0.7 mm anterior to bregma. Blue: DAPI. Red: DiI deposited on electrodes before insertion.

**FIG. S3:**
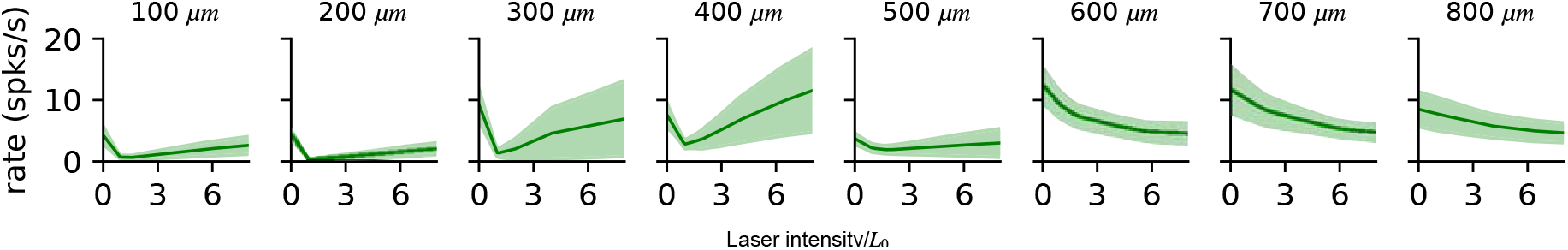
*Assoc. with Fig. 2*. Unit response to light as a function of depth. Each plot shows the average of all inhibitory units recorded from V1 in VGAT-ChR2 experiments (data as in Fig. 2) without blockers present. Panels show the average unit firing rate as a function of laser intensity at sites of different depths. The recording probes have 8 sites per shank (four shanks, 32 total sites) with 100 *μ*m spacing between sites. Inhibitory neurons on the shallowest four sites (left) show strong initial suppression to laser stimulation, and at higher laser power show increases in firing rate to stimulation. Neurons from deeper sites show suppression, but do not show a clear transition to increased firing rates, even at maximum laser intensity. This is consistent with the fact that the laser intensity experienced by neurons decays with depth due to scattering and absorption. Note that we normalize laser powers to compare excitation power across experimental sessions; this process finds just one scaling factor (*L*_0_) for all neurons and all sites for each experimental day, and the non-normalized data are shown in Fig. S4. Errorbars: SEM across units.

**FIG. S4:**
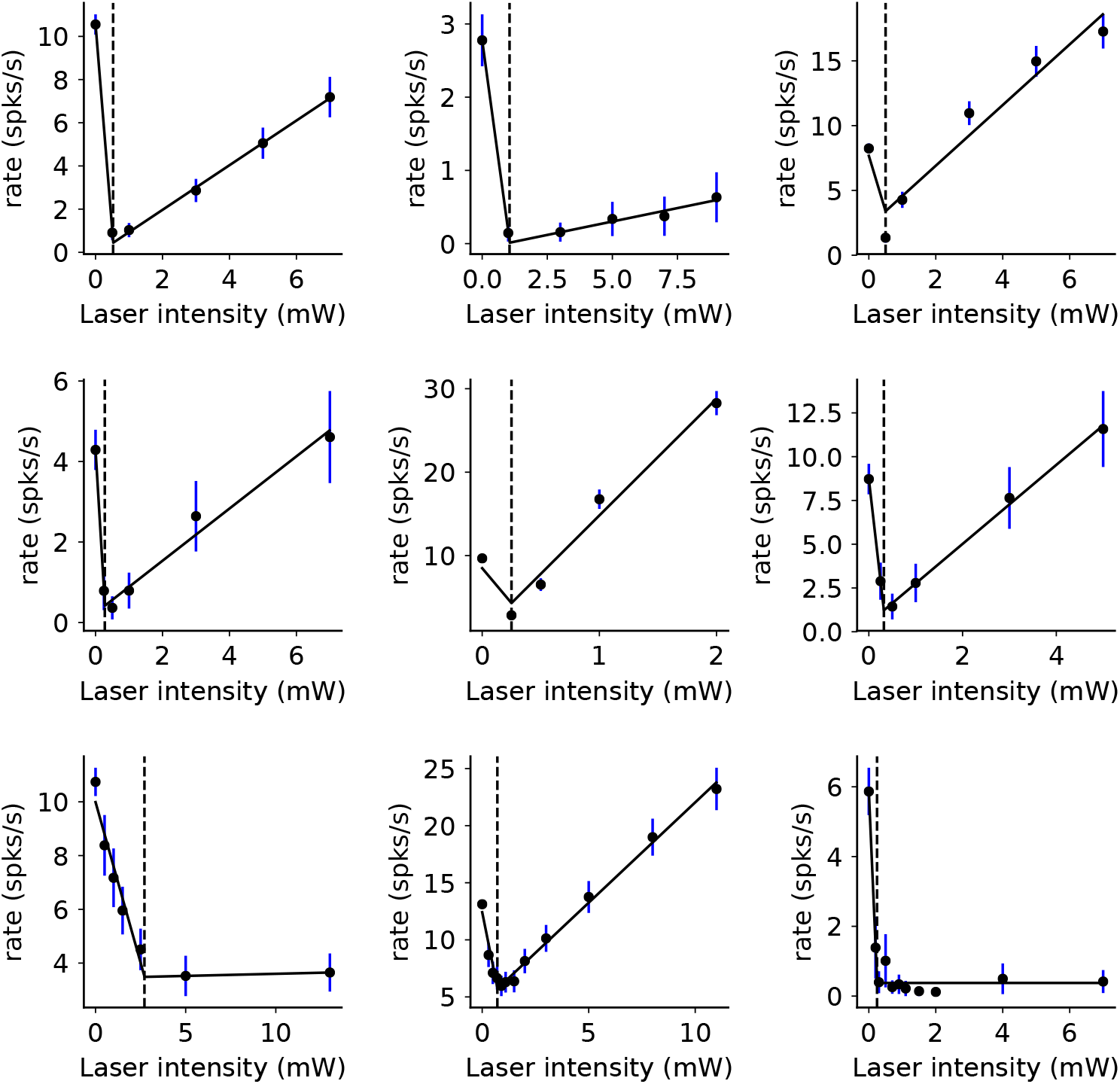
*Assoc. with Fig. 2*. Reversal point of V1 inhibitory population is typically at a low laser intensity. 9 recording sessions (panels) from V1 are shown. Dots: mean inhibitory responses (mean ±SEM) measured without blockers in the awake state. For every session, we fitted the inhibitory response to a continuous two-segment piecewise linear function (three parameters, black lines) to infer the transition point (i.e *L*_0_, black dashed lines) from ISN to non-ISN network behavior. The initial slope in all fits was negative and the second slope in all fits was non-negative (though small in three of the sessions; excluding these sessions preserved the average paradoxical effect). *L*_0_ ranged from 0.3 to 2.7 mW, presumably due to variation in the preparation such as granulation tissue thickness; we used this value to normalize intensity for combining data from different sessions. Here, as in Fig. 2, we used response with E and I blockers to classify units as E or I.

**FIG. S5:**
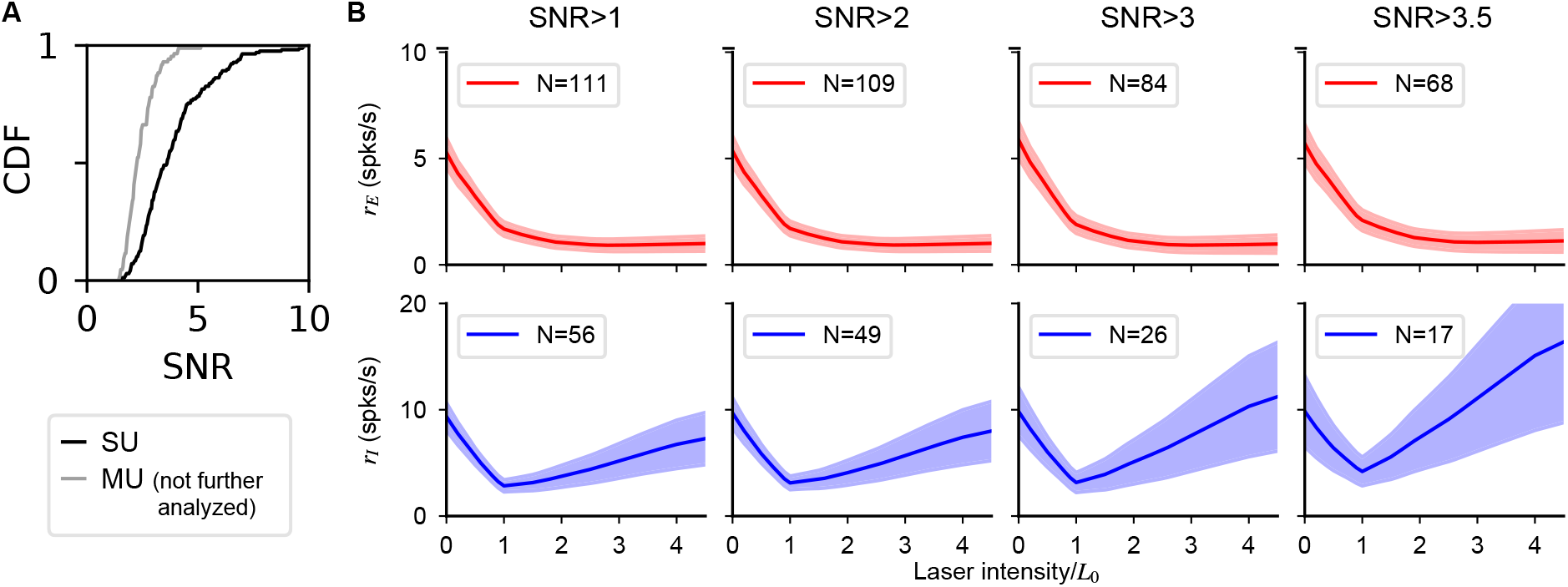
*Assoc. with Fig. 2*. Experimental results are robust to quality of unit isolation. (**A**) Signal-to-noise ratio [62] of single (black) and multi (gray) units. Single units are classified manually (blind to initial slope) based on waveform principal component separation from noise and other units, change in waveform separation over time, and inter-spike-interval distribution. As expected, classified single units have a larger SNR (single-unit E median SNR: 3.97, I median 2.83) than multi-units. Units classified as multi-units were excluded from all further analysis and are shown here only for comparison purposes. (**B**) Population response computed in a subset of SU with SNR higher than the threshold indicated in title. As the constraint becomes more stringent (left to right), the number units used decreases, and the paradoxical effect is preserved (for all panels, initial slope median is significantly negative: all *p* < 10^−5^ except bottom right: *p* < 0.003; one-sided Mann-Whitney U test). Additional evidence that the paradoxical effect is not dependent on our unit isolation comes from the fact that the PV-Chronos recordings were conducted with identical experimental procedures, and show no average paradoxical effect (see Results and Fig. 6).

**FIG. S6:**
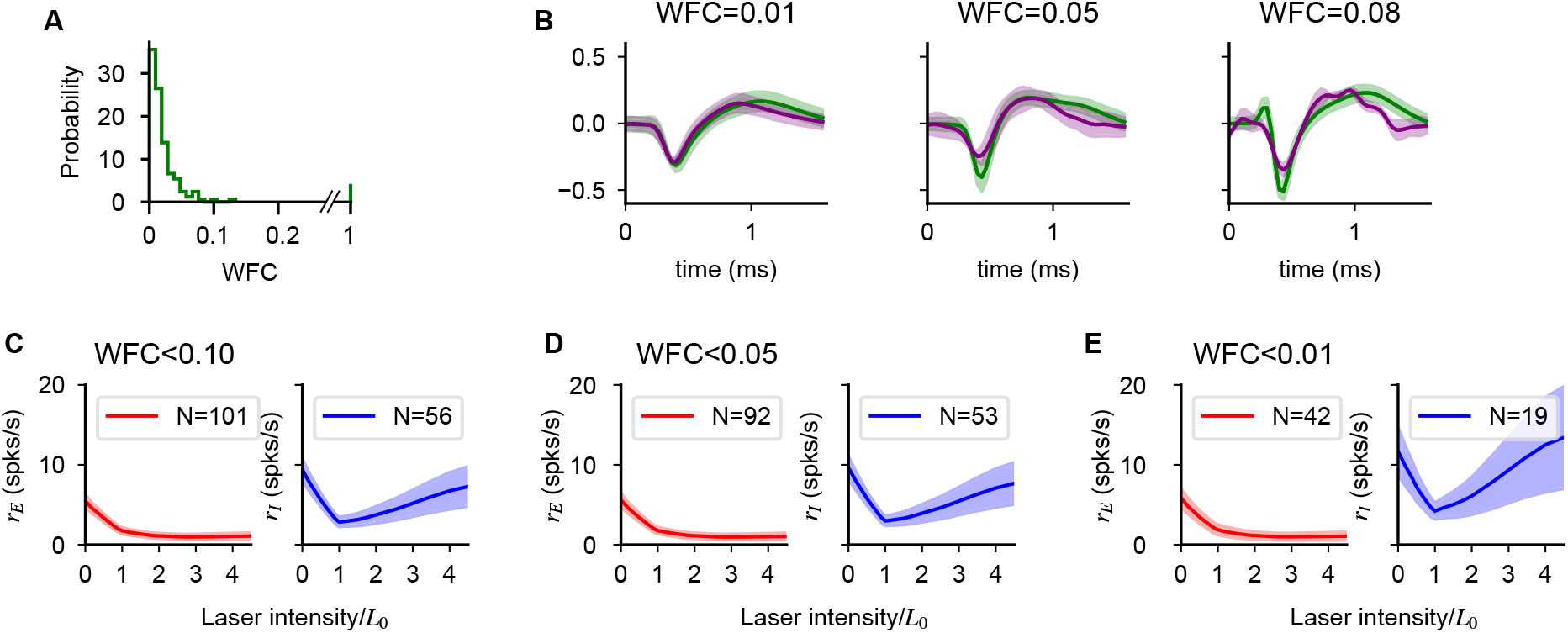
*Assoc. with Fig. 2*. Waveform drift has no qualitative effect on population response. (**A**) Distribution of relative waveform change index (WFC). WFC is defined as, for every unit, the quantity

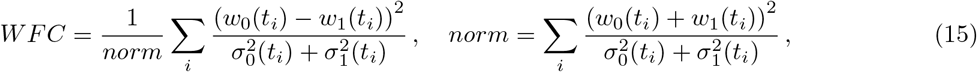

where the waveform and its variance are given by *w_j_*(*t*) and 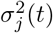, and *j* = 0,1 indexing the blocker, no-blocker cases. *t_i_* ranges over time samples of the recorded waveforms (45 samples/waveform, 1.5 ms, 30 kHz acquisition rate). Eq. (15) measures the relative change in waveform of a unit before and after the addition of blockers. Note that a small number of units did not generate any spike after the addition of blockers (WFC=1, N=9); these could either be excitatory units that become silent, or units that drifted substantially during the recording. (**B**) Example single unit before (green) and after (purple) the addition of synaptic blockers. (**C-E**) Population response of excitatory and inhibitory cells computed using single unit selected with a threshold on the maximum waveform change. As the constraint becomes more stringent (left to right), the number units used decreases but the structure of the response is not modified and the paradoxical effect is preserved. Our waveform-sorting procedure uses a contour, fixed over time, in the 2d axis of the first two principal components (PCs) to classify units. Therefore, larger WFC values signify shifts in waveform (e.g. WFC=0.08 example, B, right panel), despite the fact the limits of maximum waveform shift are constrained to a range of possible PC values specified by the contour. Including units that show relatively larger waveform shifts (compare D and E) does not modify our effects.

**FIG. S7:**
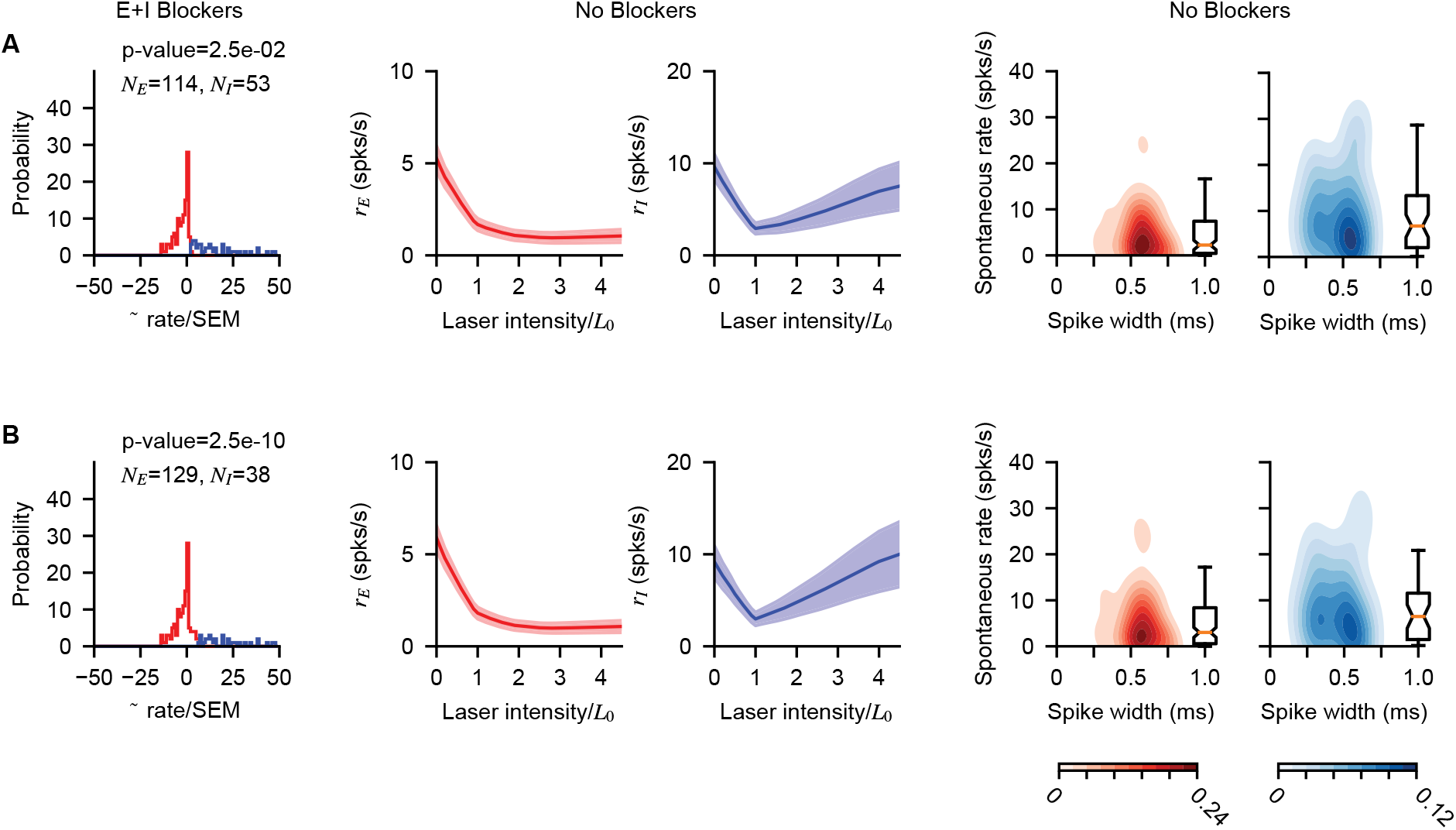
*Assoc. with Fig. 2*. Paradoxical V1 inhibitory response does not depend on statistical E/I classification threshold. The pharmacological classification method (based on firing rate increase to optogenetic stimulation when E+I blockers had been applied; Fig. 2) is robust to choice of classification threshold applied to that firing rate change. (In addition, Supp Fig. S5 shows paradoxical effect is preserved as waveform isolation via SNR is varied; Fig. S6 shows potential waveform drift over time does not affect results.) (**A**) Left column, distribution of response slopes shown with a permissive classification threshold (*p* < 0.05; to make the test two-tailed we used *p* < 2.5 · 10^−2^; Welch’s t-test allowing unequal variances). Middle two columns: resulting average response to stimulation. Right two columns: distribution of spontaneous rates and waveform widths. ((**B**)) Same, but for a more restrictive classification threshold (*p* < 10^−9^). Decreasing the threshold reduces the number of units classified as inhibitory, (13 units re-classified as excitatory once the more stringent threshold is applied), but does not modify the conclusions of the main text; in particular paradoxical inhibitory modulation is preserved.

**FIG. S8:**
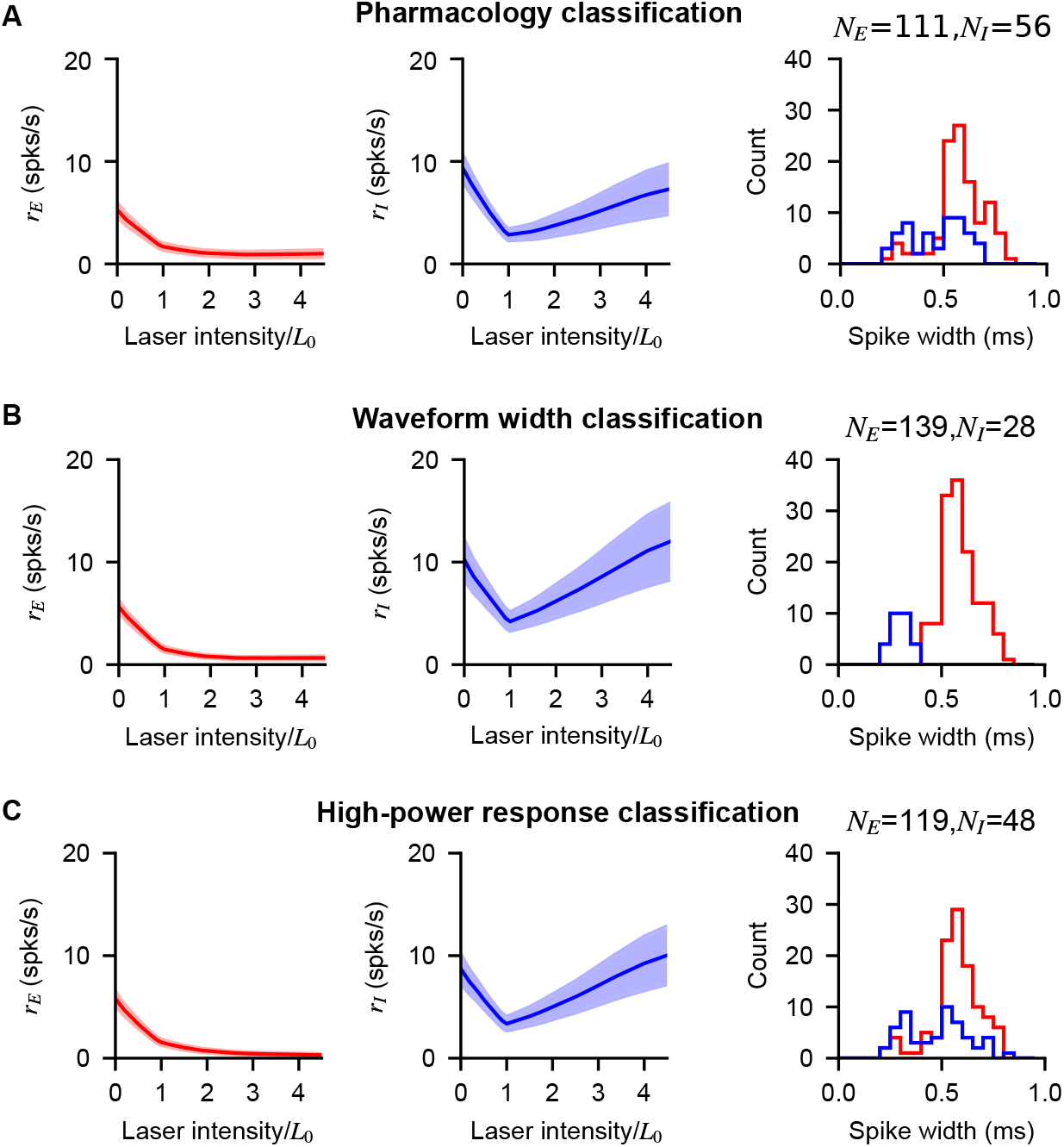
*Assoc. with Fig. 2*. Paradoxical effects in V1 are preserved when inhibitory units are classified via pharmacology, via waveform width, and via response at high laser power. Population response in V1 of excitatory and inhibitory cells as defined by: response with blockers (**A**), waveform width (**B**), and firing rate increase at high laser intensity (**C**). Fig. 2 of the main text uses the classification in shown in (A), but a similar population response results using the two alternative methods of classification. (**B**) units are classified as inhibitory if their waveform width is less than 0.45 ms, a value determined by the bimodal distribution observed in the population (left panel). (**C**) units are classified as inhibitory if their firing at maximum laser intensity (5*L*/*L*_0_ in the figure) is one SEM larger than their firing at *L*_0_ (the minimum inhibitory network response).

**FIG. S9:**
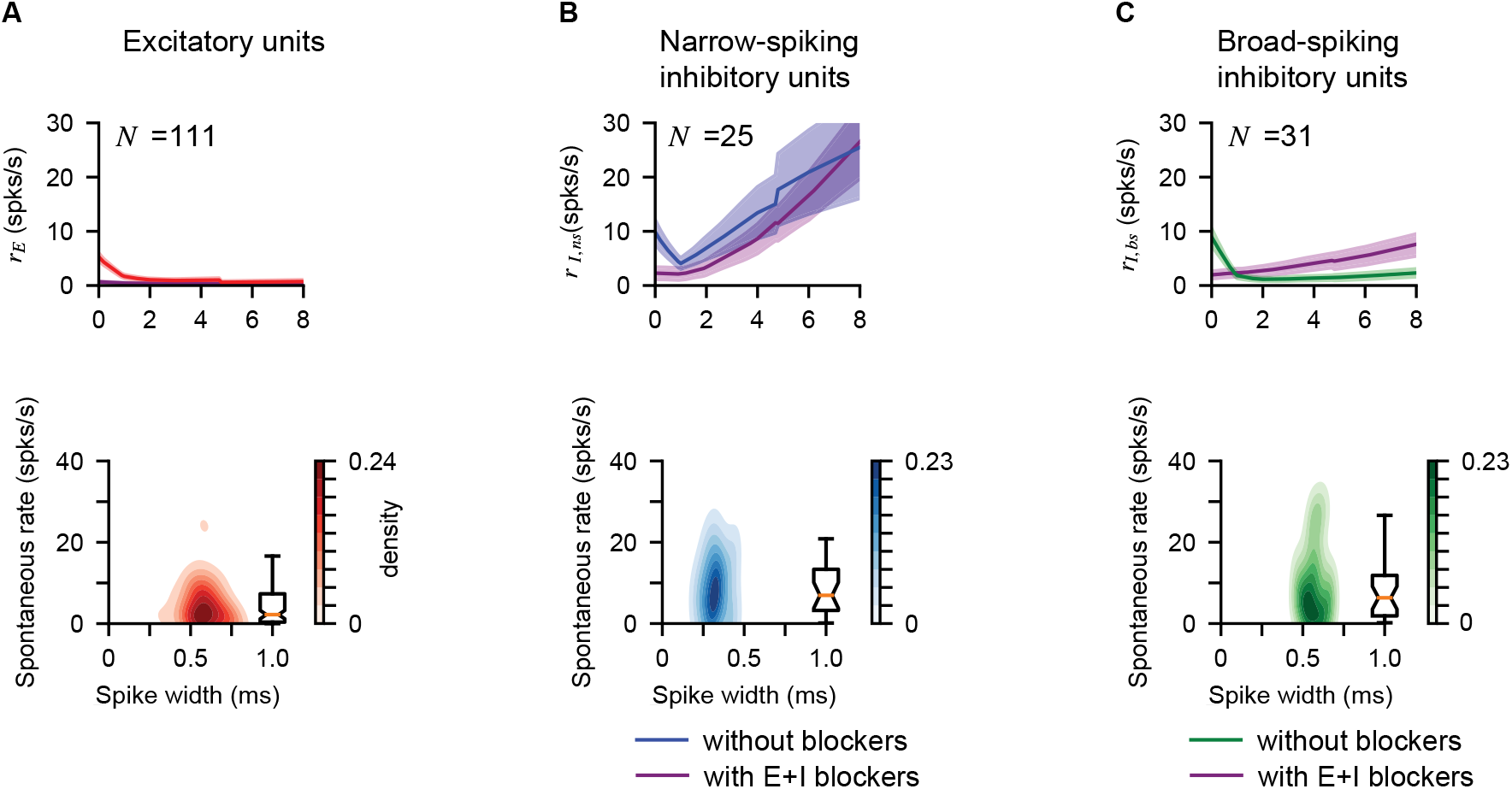
*Assoc. with Fig. 2*. Responses of broad- and narrow-spiking V1 (VGAT-ChR2) inhibitory units. Average rate (first row) and distribution of spontaneous rates vs spike width (second row) of excitatory units (**A**, *N* = 112), narrow-spiking inhibitory units (**B**, waveform width < 0.45ms, *N* = 24), and broad-spiking inhibitory units (**C**, waveform width> 0.45ms, *N* = 30). Corresponding average response with E+I blockers is plotted to show elevated responses with light (first row, purple). Cells are classified as E or I via pharmacological classification (as in Fig. 2). The two inhibitory populations both show initial paradoxical suppression. Narrow-spiking units show a more pronounced increase in firing rate in the non-ISN (high laser power) regime. This difference seems unlikely to result from experimental differences like variation in opsin expression levels, and more likely may be generated by recurrent inhibition. (*i.e*. narrow spiking putative basket/PV cells provide strong inhibition to wider-spiking somatostatin or other inhibitory cells, and at high laser powers PV cells suppress the wider-spiking units.) In Fig. 6, for both the viral PV-Chronos and transgenic PV-ReachR experiments, we classify cells as inhibitory based on elevated response at high laser power. Consistent with the V1 VGAT-ChR2 data shown above, the cells in PV experiments classfied as inhibitory by their high-power response also have narrow waveforms on average.

**FIG. S10:**
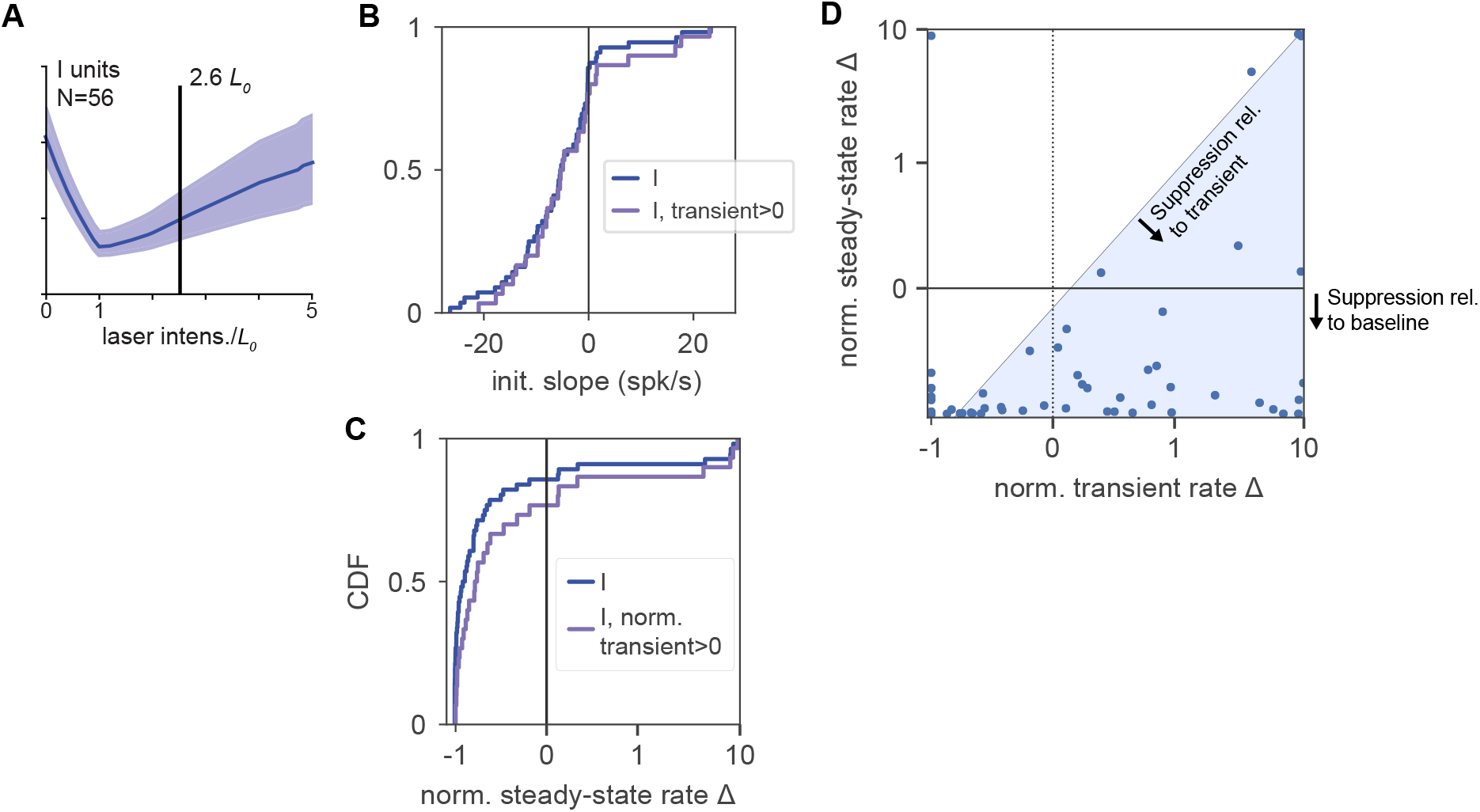
*Assoc. with Fig. 3*. Relationship between inhibitory initial transient and paradoxical suppression. (**A**) Schematic of laser intensity used here and in Fig. 3. This intensity (2.6 · *L*_0_) is above the point of maximal paradoxical steady-state suppression (1.0 · *L*_0_), but shows the initial transient (Fig. 3AB) clearly. (**B**) Paradoxical suppression (initial slopes < 0) is seen even when selecting only the inhibitory units that show a positive initial transient (“transient > 0”), those units with a an increase in spike rate over an interval [1,12] ms after light onset. (“I”: all recorded inhibitory units, N=56). (**C**) Paradoxical suppression is preserved for transient-only units: same as (B), plotting the distribution of steady-state responses. Normalized responses < 0, to the left of the vertical line at 0, are units that are suppressed (i.e. that show paradoxical suppression) in steady state, after the end of the transient, for this laser intensity. (**D**) The majority of recorded inhibitory units show steady-state paradoxical suppression relative to both baseline rate and to transient rate. y-axis: steady-state rate change (normalized to baseline rate; i.e. 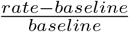, a value < 0, labeled at right, signifies paradoxical suppression). x-axis: transient rate change (normalized to baseline rate; I units with an initial positive transient have values > 0). The majority of the units are in the lower right quadrant (transient > 0, steady-state < 0). A few units show a positive transient and elevated steady-state rate (upper right quadrant); these are likely inhibitory units showing non-paradoxical steady-state rate increases. Blue shaded region shows units whose normalized steady-state rate is lower than their normalized transient rate. The fact that most inhibitory units fall into this region shows that steady-state suppression does not arise from mixing E and I waveforms, but instead that we accurately record inhibitory units — which often show initial transients followed by suppression as predicted by ISN models.

**FIG. S11:**
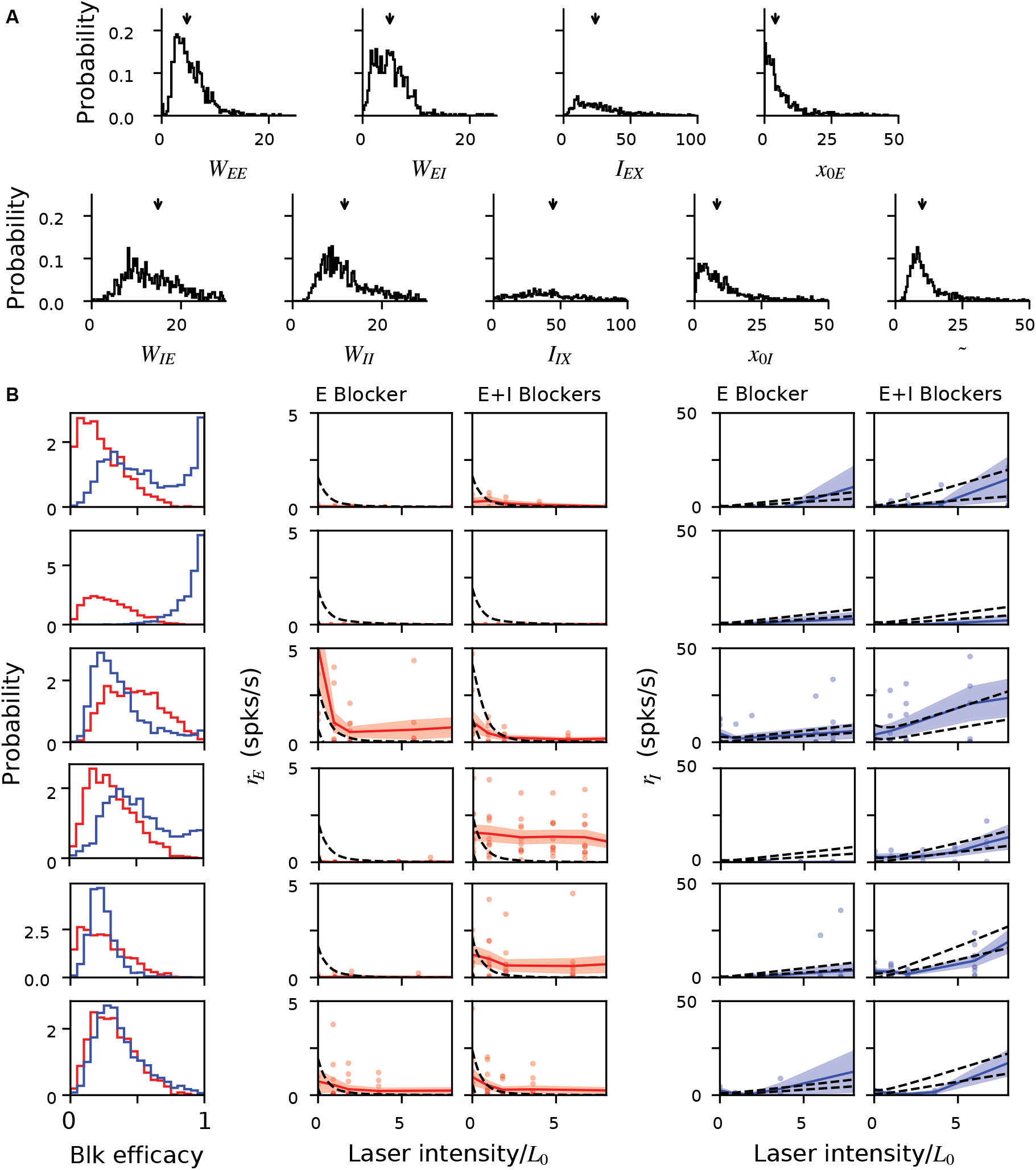
*Assoc. with Fig. 4*. V1 model parameter stability shown via data bootstrap. To examine the stability of the model fit shown in Fig. 4, here we show the distribution of the parameter estimates obtained via bootstrapping (randomly resampling) the data (Methods). (**A**) Distribution (solid lines) of the network parameters (defined by Eq. (6)), and corresponding medians (arrows) over all the bootstrap runs. (**B**) Distribution of the parameters for excitatory (red) and inhibitory (blue) blocker efficacy (left column) with corresponding measured and predicted responses (middle column, red: excitatory cells; right column, blue: inhibitory cells). Different rows correspond to different recording sessions. Shaded areas: ±1 SEM across units. The efficacy of blockers varies somewhat from day to day, justifying our use of different efficacy parameters for different recording days in the model.

**FIG. S12:**
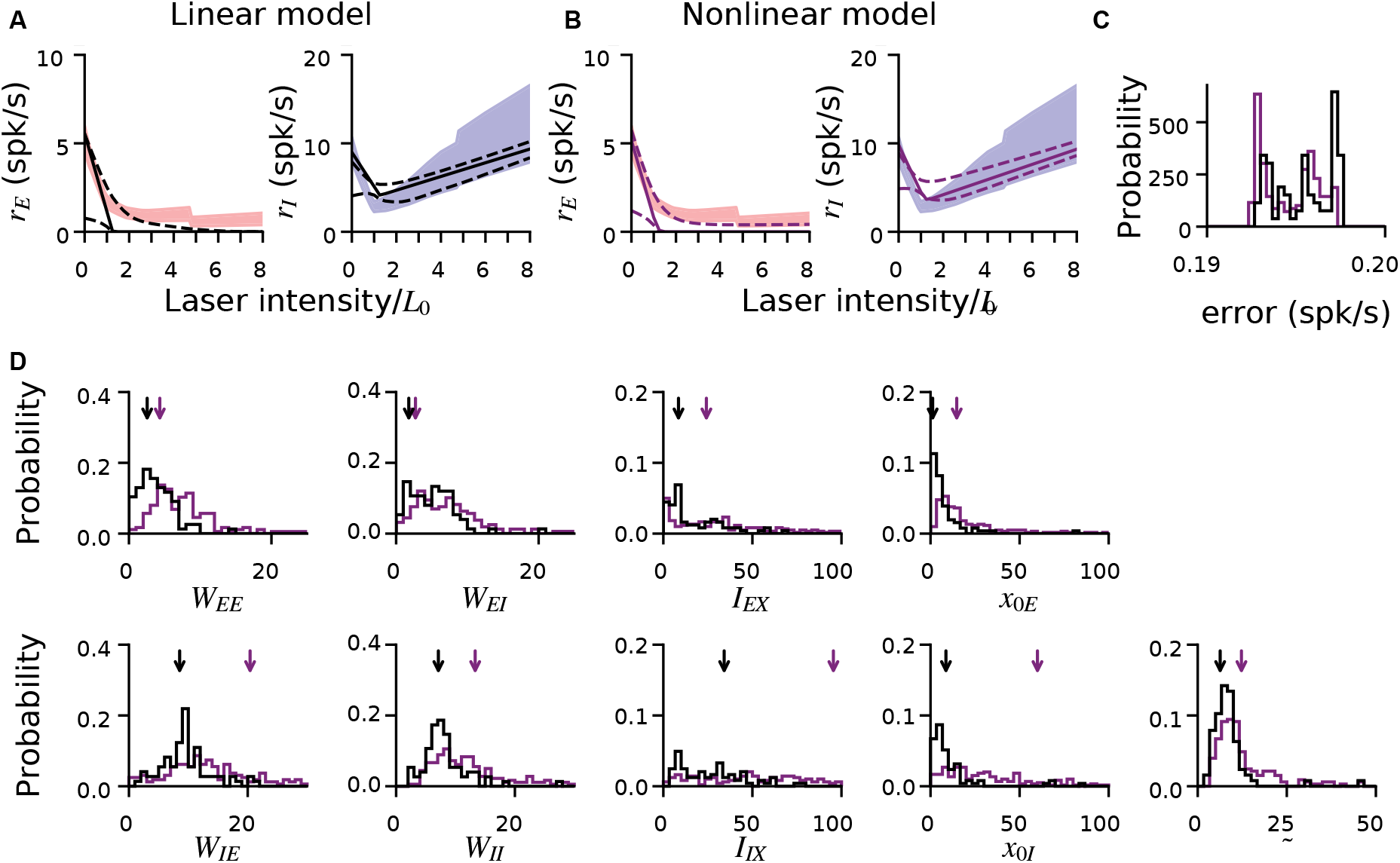
*Assoc. with Fig. 4*. Comparison of linear and nonlinear rate models, and global optimization method. (**A-B**) Similar characterizations of the V1 data are provided by linear and non-linear rate models (i.e. models with either piecewise-linear or non-linear single-cell transfer functions; Methods), even though the non-linear model has one extra parameter (which controls the transfer function nonlinearity; Methods). Shaded regions: means ±1 SEM for E (red) and I (blue) units; data is recorded from V1 of awake animals, same data set as Figs. 2–4. Solid black lines: optimal (lowest-error) fit, across many optimization runs initialized with different parameter values. The lowest-error fit is the global minimum of the optimization cost (error) function. To include only plausible solutions, we retained optimzation results only where error is within 2.5% of optimal. Dashed lines: ±1 s.d. across optimization runs. (**C**) Error values for the two models (black: linear; purple: nonlinear). Error (units: spk/s) is the absolute difference between model and data mean, averaged over all laser intensities. (**D**) Parameters corresponding to solution shown in A,B. While the optimal solutions (arrows) show some differences between the two models, the distribution of parameters across runs overlaps substantially, showing the inferred network parameters are robust to the specific model used. Note that the distributions shown here are taken over optimization initialization values, in contrast to Fig. 4, which shows variability of the optimal solution over bootstrap runs.

**FIG. S13:**
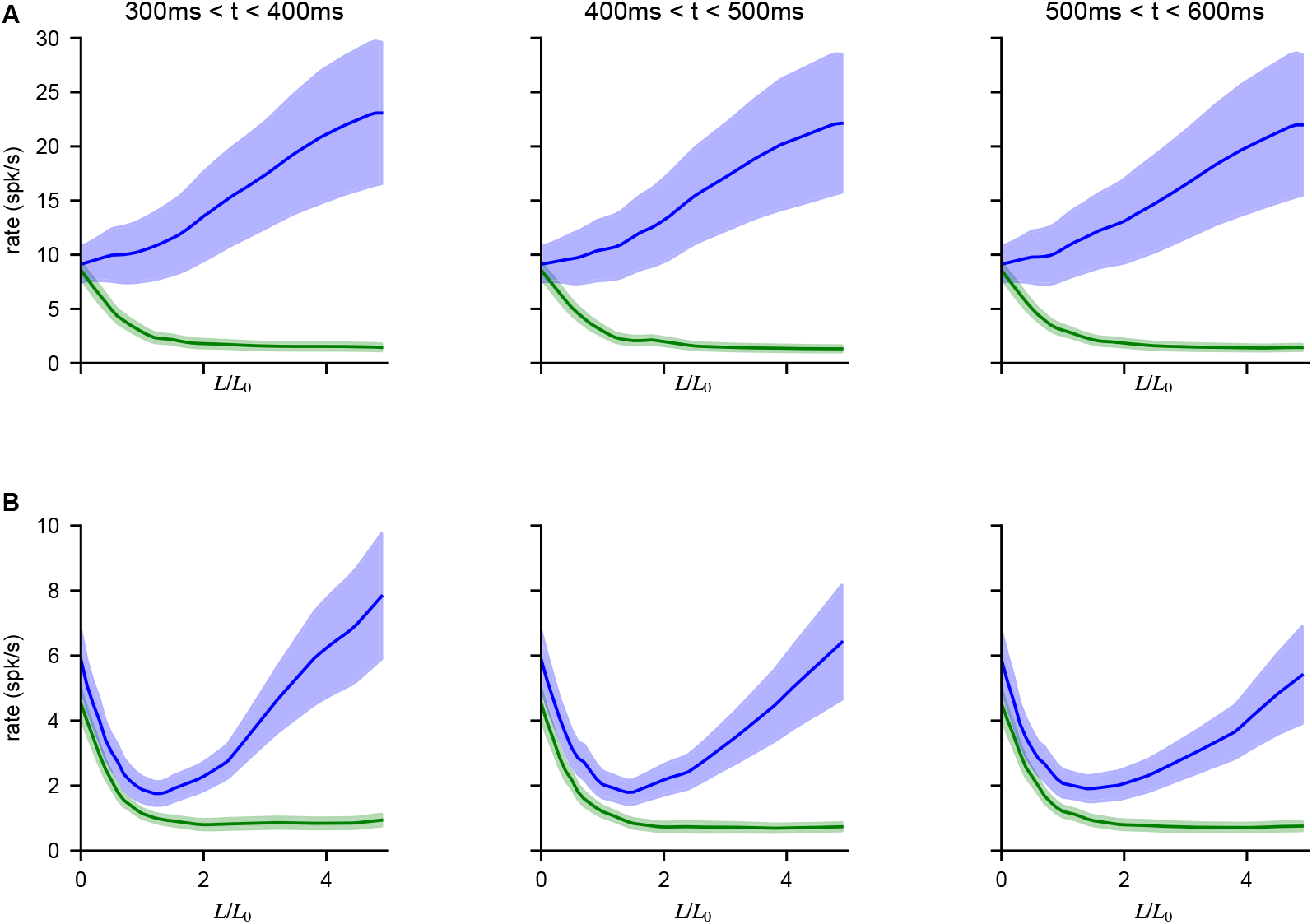
*Assoc with Fig. 6*. No effect of opsin kinetics on paradoxical effects. (**A**) Average spike rate responses from PV-Chronos stimulation over different counting windows (300-400 ms, 400-500 ms, 500-600 ms). The overall effect is unchanged, showing that we are measuring steady-state network response, and onset and offset opsin kinetics does not affect our paradoxical effect measurements. (**B**) same as A, but for PV-ReaChR recordings. Units shown here are the same as in Fig. 6d,f, where the firing rates are found over the whole (300-600 ms) counting window.

**FIG. S14:**
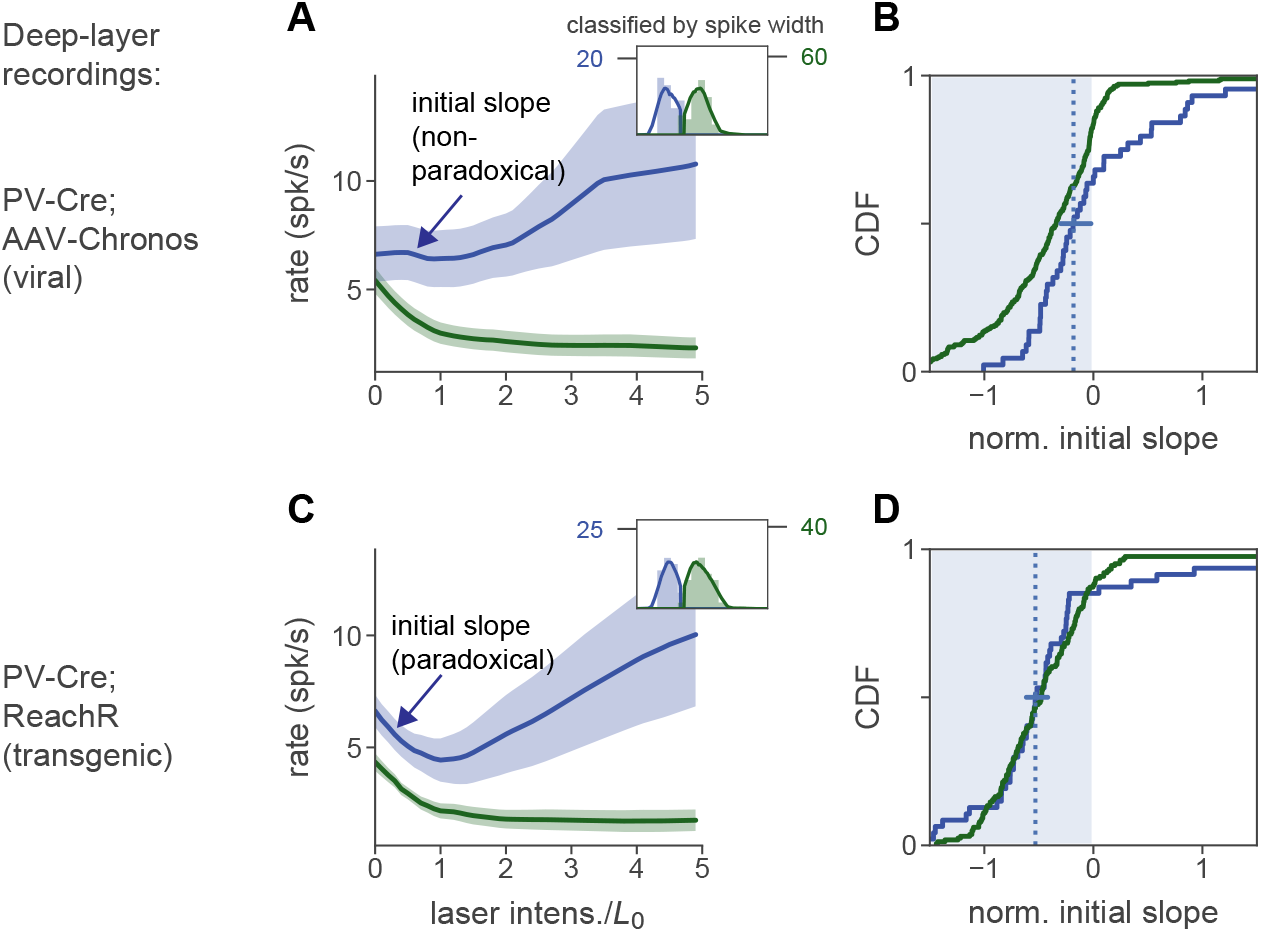
*Assoc. with Fig. 7*. Effects of PV stim with viral vs. transgenic expression in deep-layer neurons is similar to effects in superficial layers. (**A**) Deep-layer (depth ≥ 500 *μ*m) units from viral transfection experiments yield no mean paradoxical effect, as in superficial-layer recordings (Fig. 7). (**B**): initial slopes for all recorded units: both mean and median are negative (horiz line, 95% CI for median via bootstrap). (**C-D**) Deep-layer units with transgenic expression do show a mean paradoxical effect. Conventions as in Fig. 7; blue indicates inhibitory cells, and green indicates excitatory plus non-PV plus PV non-transfected neurons. As in Fig. 7, these neurons are classified by waveform width. In the transgenic line (C-D), paradoxical suppression is clear, but less strong in deep layers than in superficial layers (*cf* Fig. 6), a pattern we also saw with stimulation of all inhibitory cells (Figs. 2,5,7). Because PV-ReaChR was stimulated with red light, we expect more direct effects of stimulation in deeper layers than with blue-light VGAT-ChR2 stimulation (Fig. 7), and indeed, here a greater number of inhibitory cells show paradoxical suppression (D) than in the VGAT-ChR2 deep layer recordings from V1 (Fig. 7d). Note that because we stimulate PV neurons and not all inhibitory cells, the data in this figure is not diagnostic of ISN operation, but illustrates that PV stimulation can indeed produce paradoxical effects in inhibitory neurons in deep layers.

**FIG. S15:**
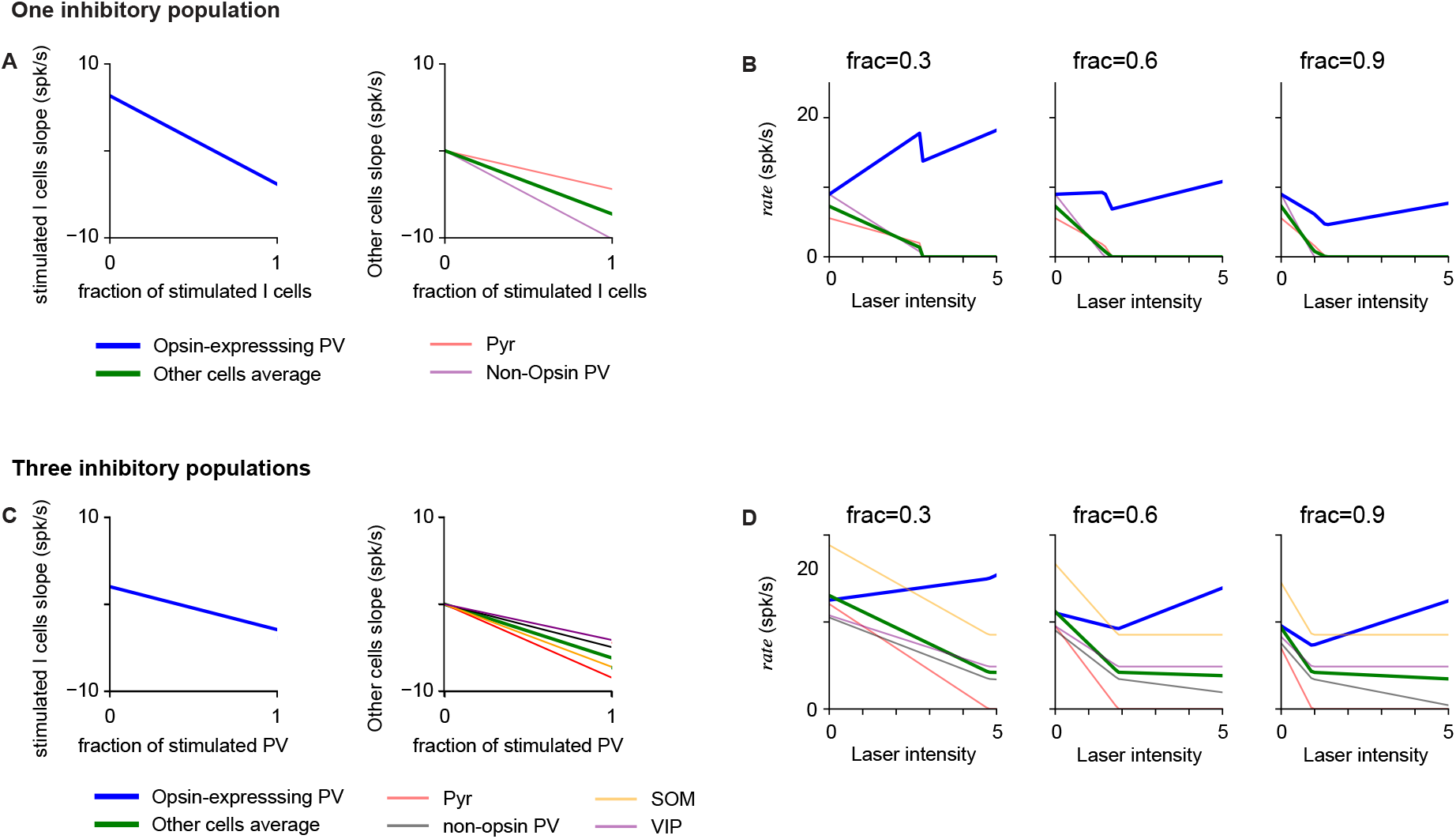
*Assoc. with Fig. 6*. Additional analysis of inhibitory responses to partial stimulation of inhibitory population. (**A-B**) Network response computed from Eq. (10) for different fractions of stimulated inhibitory cells. (**A**) Response slope at baseline activity of different populations in the network. The response slope of stimulated cells is positive when the fraction is small and becomes positive as the fraction increases. For these model parameters (computed from data, Fig. 4), the response slope of the non-stimulated cells is negative for all fractions. (**B**) Responses as a function of laser intensity for different fractions of stimulated cells (title). (**C-D**) Results analogous to those in A-B, but obtained in a model with three inhibitory populations (representing PV, SOM, VIP [19]) as a function of the fraction of stimulated PV cells (i.e. opsin-expressing PV cells). The model structure is analogous to that of Eq. (10) but with three inhibitory populations, one of which is partially stimulated. The connectivity matrix used is consistent with experimental measurements [39] and is given by: *W_AE_* = 1.2 if *A* = *E* and 1 if *A* = *PV, SOM, VIP*; *W_APV_* = −1 if *A* = *E, PV* and 0 if *A* = *SOM, VIP*; *W_ASOM_* = −1 if *A* = *E*, −0.5 if *A* = *PV*, 0 if *A* = *SOM* and −0.6 if *A* = *VIP*; *W_AVIP_* = 0 if *A* = *E, PV, VIP* and −0.25 if *A* = *SOM*.

**FIG. S16:**
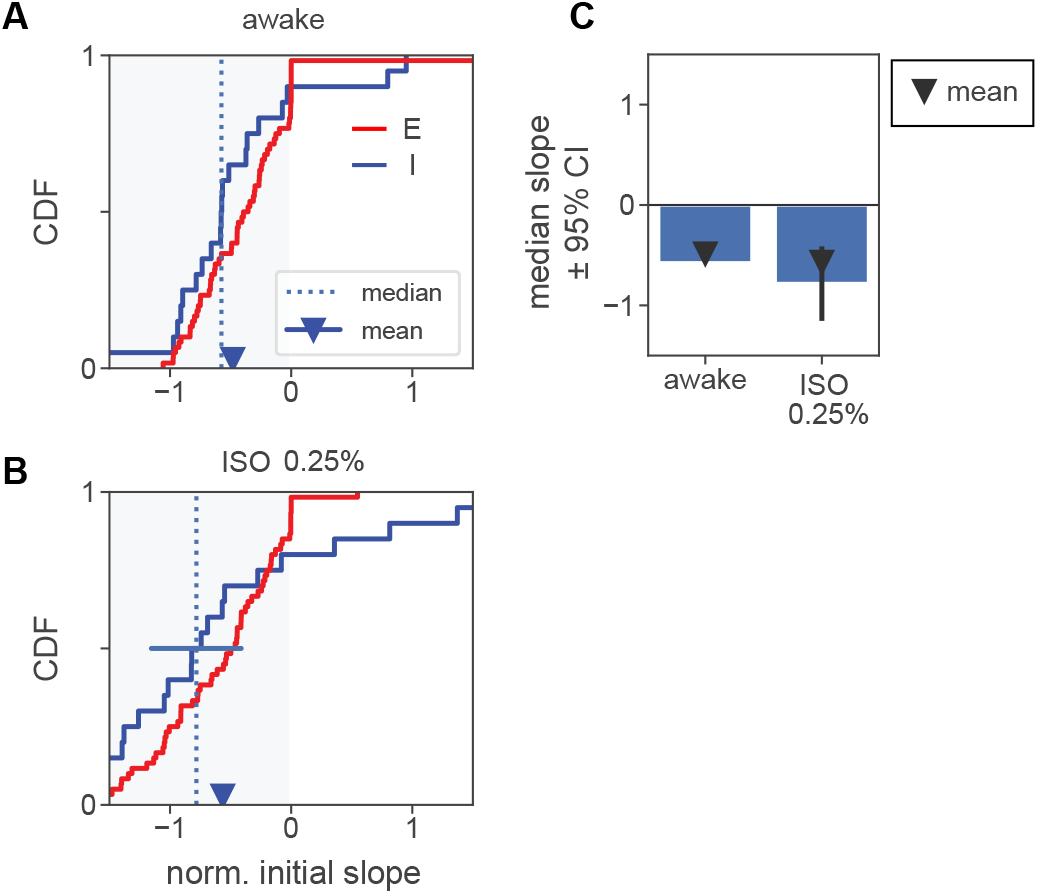
*Assoc. with Fig. 8*. Normalized initial slopes for units recorded in anesthesia experiments. (**A**) Normalized slopes for E and I units in awake recordings. Conventions as in Fig. 5d. (**B**) Normalized slopes for E and I units in recordings with 0.25% isoflurane. More inhibitory units have a positive initial slope than in (A), but the mean and median remain negative. This increase in heterogeneity may be due to the network moving closer to an ISN transition point as firing rates are decreased by anesthesia. (**C**) Summary of mean and median slopes for inhibitory units. Both mean and median are significantly negative in both cases. (Mean, awake: *p* < 0.01, iso: *p* < 0.05, via t-test; median, awake: *p* < 10^−5^, iso: *p* < 10^−3^, Mann-Whitney U.) Solid black bars: 95% confidence interval for median via bootstrap.

## Appendix A: Response analysis with networks of leaky integrate and fire neurons

In the main text, using rate models without input noise, we have shown that experimental data recorded in V1 are consistent with the network operating in the inhibition stabilized regime, with mean excitatory and inhibitory inputs larger than threshold and total input of order threshold. In this section we show that similar results are obtained using a rate model which features input noise and a more biologically accurate transfer function. Furthermore, this analysis shows that inferred parameters agree with previously measured network properties, and that data are consistent with the loose balance regime [33] and noise driven firing.

### Model definition

We use a network of excitatory and inhibitory neurons analogous to the one described in the main text, but with a single neuron transfer function matching that of leaky integrate and fire models. In the network, each neuron in population *A* = [*E, I*] receives a Gaussian distributed input, of mean *μ_A_* and variance 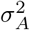, which is produced by recurrent and feedforward interactions. As showed in [63], population firing rates are found solving:

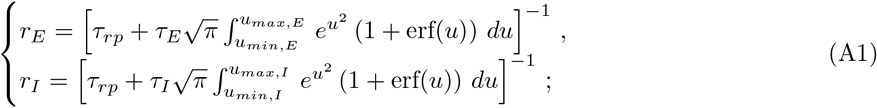

where *τ_rp_* is the single neuron refractory period, while *u_max,A_* and *u_min,A_* are the distance form spiking threshold *θ* and reset *V_r_* of the mean input *μ_A_* measured in units of input noise *σ_A_*, i.e.

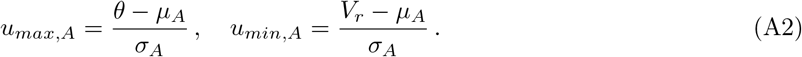

In the model, using the same notation as in the linear model of the main text, means are given by

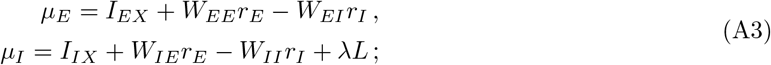

we assume noise amplitude to be fixed to *σ_E_* = *σ_I_* = 5 mV [64]. Other model parameters are: *τ_rp_* = 2ms, *τ_E_* = 20ms, *τ_I_* = 10ms, *θ* = 20mV, *V_r_* = 10mV. This model has a total of 9 free parameters (4 connectivity parameters *W_AB_*, 2 feed-forward inputs *I_AX_*, 1 laser efficacy λ, 2 blockers efficacy *ϵ_A_*) which will be inferred from the data.

### Fitting procedure

Evaluating rates from Eq. (A1) is computationally expensive; to speed up the exploration of the parameter space, we fitted the model directly to the average population response as follows.

We discretized laser intensities into 101 bins centered around laser values *L_i_* and equally spaced in the interval [0, *L_max_*] (*L_max_* is the maximum *L*/*L*_0_ common to all recording days, we used *L_max_* = 15 and checked that results were unchanged for *L_max_* = 10–14). The error in the model description was computed as

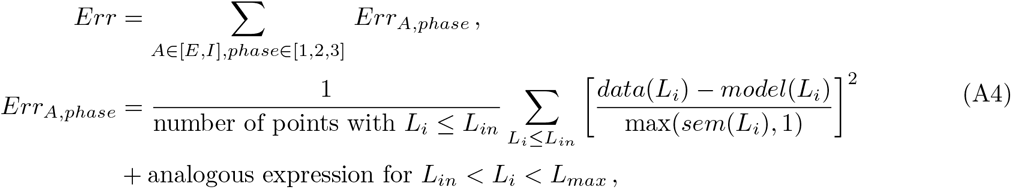

where *model*(*L_i_*) is the model prediction obtained from Eq. (A1) for *L* = *L_i_*, while *data*(*L_i_*) (*sem*(*L_i_*)) is the mean rate (standard error of the mean); this was computed averaging responses from all cells to stimuli of intensity *L*/*L*_0_ within the bin centered around *L_i_*. The parameter *L_in_* divides data into two groups, *L_i_* < *L_in_* and *L_in_* < *L_i_*, whose contributions to the error are weighed by the corresponding number of data points; this split was used to prevent the large laser responses of inhibitory cells from dominating the error (we used *L_in_* = 1 and checked that results were robust for *L_in_* = 2, 3). The minimum bound on the standard error of the mean (max(*sem*(*L_i_*), 1)) prevents regions with low *sem*(*L_i_*) (specifically, the large laser intensity response of excitatory cells) from dominatint the error.

Given the average neural responses of different populations and phases, optimal model parameters were determined by minimizing Eq. (A4) as described in the methods section of the main text. Finally, to better estimate the excitatory rate at large laser intensity, we included in our analysis only cells that were classified in the same way (inhibitory or excitatory) from the classification done with blockers and with response at high laser intensity. This filtering reduces the number of cells used in the analysis, from 167 (111 E+ 56 I) to 119 (90 E + 29 I), and decreases the excitatory rate measured at large laser intensity (Fig. S17A).

### Results of the analysis

The best model description of the data, obtained minimizing Eq. (A4), is shown in Fig. S17A. The corresponding model parameters are:

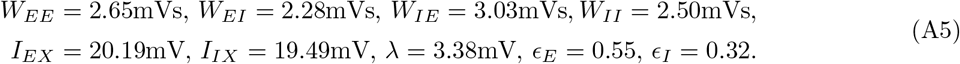

As in the main text, we used a bootstrap approach to estimate the precision with which model parameters can be inferred from the data; results are shown in Fig. S17B. The analysis shows that distribution of inferred parameters are localized around the optimal solution described above. Moreover, the parameters of the model can be map to known biophysical quantities by noticing that *W_EE_* = *τ_E_K_EE_J_EE_* [63], where *K_EE_* and *J_EE_* are the average number of recurrent excitatory projection and their efficacy (analogous expression holds for the other elements of the matrix *W*). Assuming *τ_E_* = 20ms and *K_EE_* ≈ 10^2^ – 10^3^, we obtain an estimate of *J_EE_* ≈ 0.1 – 1mV, which is consistent with direct biophysical measurements [65].

We computed the inputs into excitatory cells; contributions coming from recurrent inhibition (−*W_EI_r_I_*), recurrent excitation (*W_EE_r_E_*), and feed-forward excitation (*I_EX_*) are shown in Fig. S17C. The total input is found to be below threshold, at a distance of order one in unit of input noise. This result is consistent with the balanced state model [5; 6; 7], where firing is driven by input noise. However, unlike what is expected by the balanced state model, the different components of the input (e.g. feed-forward excitation) are of order threshold, consistent with the loose balance regime recently suggested to underlie cortical dynamics [33].

In the model, the strength of self-excitation of the excitatory population is obtained from Eq. (A1) as

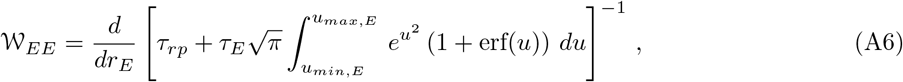

where the integral is evaluated at the inferred baseline activity. The quantity 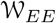 is the generalization of the parameter *W_EE_* analyzed in the main text, which takes into account the amplification of coupling strength due to transfer function nonlinearities. Values of 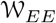 inferred with data bootstrap are shown in Fig. S17D. Results are consistent with 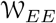 larger than one, indicating that indeed the excitatory subnetwork is unstable without inhibition, and in the range 5-15, consistent with the result obtained in the main text using a linear model.

To summarize, the analysis derived in this section shows that results obtained in the main text are preserved when a more biologically accurate, spiking, model is used to describe the data. Moreover, this approach also shows that inferred parameters are consistent with known biophysical properties of cortical networks.

**FIG. S17:**
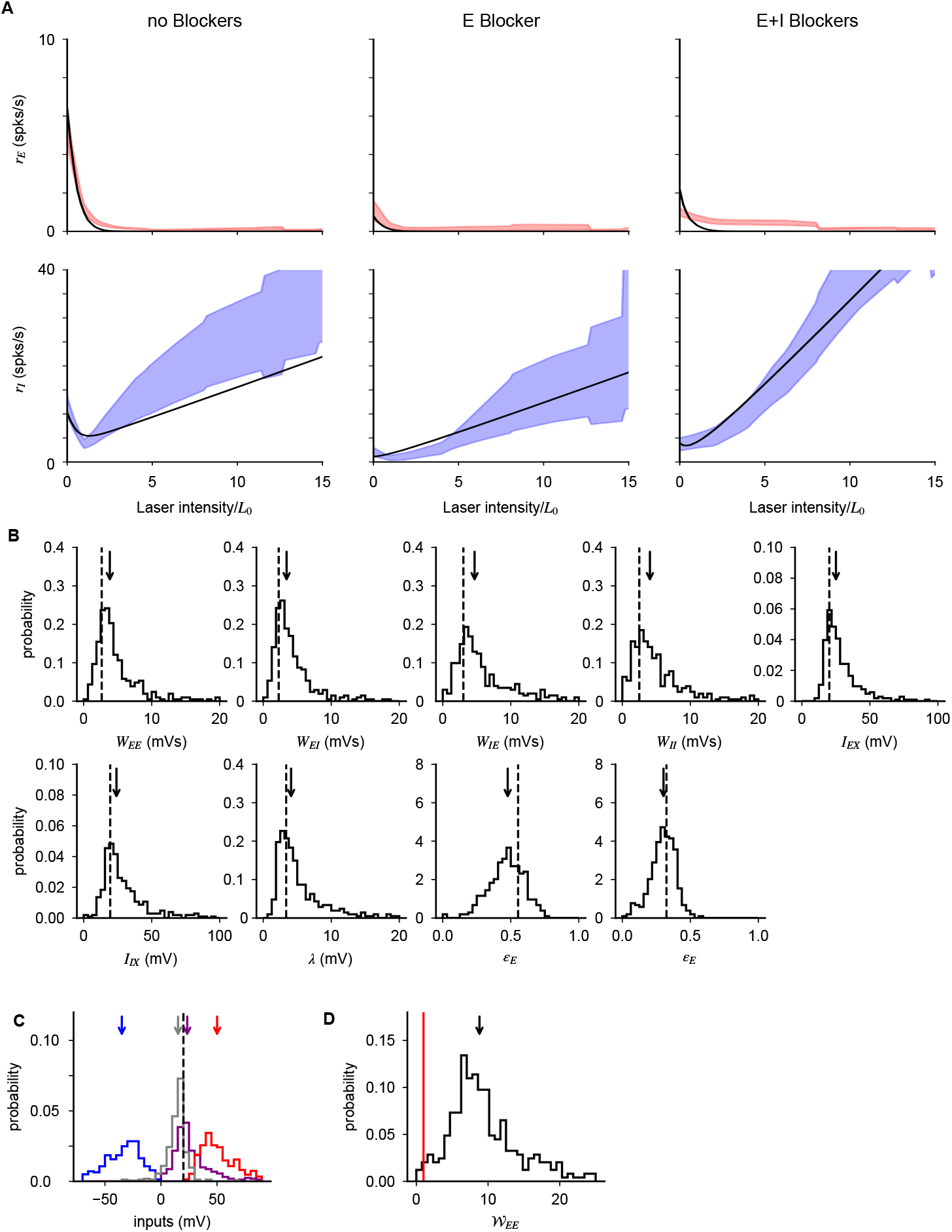
Population response in the three phases of the experiment. (**A**) Optimal fit of population response given by Eq. (A1). (**B**) Distributions (lines) and medians (arrows) of inferred parameters obtained with data bootstrap; dashed lines are optimal parameters reported in the text. (**C**) Distribution (line) and median (arrow) of self excitation of excitatory cells computed from data using Eq. (A6). The inferred values are distributed above the instability point *W_EE_* = 1 (red line). (**D**) Distributions (lines) and medians (arrows) of recurrent excitation (*W_EE_r_E_*, purple), recurrent inhibition (−*W_EI_r_I_*, blue), feed-forward+recurrent excitation (*W_EE_r_E_* + *I_EX_*, red), and total input (−*W_EI_r_I_* + *W_EE_r_E_* + *I_EX_*, gray) to excitatory cells. As discussed in the text, inputs are of order threshold (dashed line); data are consistent with fluctuation driven firing [5; 6; 7] and loose balance regime [33].

